# Nix induced mitochondrial fission, mitophagy, and myocyte insulin resistance are abrogated by PKA phosphorylation

**DOI:** 10.1101/825828

**Authors:** Simone Cristina da Silva Rosa, Matthew D. Martens, Jared T. Field, Lucas Nguyen, Stephanie M. Kereliuk, Yan Hai, Donald Chapman, William Diehl-Jones, Michel Aliani, Adrian R. West, James Thliveris, Saeid Ghavami, Christof Rampitsch, Vernon W. Dolinsky, Joseph W. Gordon

## Abstract

Lipotoxicity is a form of cellular stress caused by the accumulation of lipids resulting in mitochondrial dysfunction and insulin resistance in muscle. Previously, we demonstrated that the mitophagy receptor Nix is responsive to lipotoxicity and accumulates in response to diacylglycerols induced by high-fat (HF) feeding. In addition, previous studies have implicated autophagy and mitophagy in muscle insulin sensitivity. To provide a better understanding of these observations, we undertook gene expression array and shot-gun metabolomics studies in soleus muscle from rodents on an HF diet. Interestingly, we observed a modest reduction in several autophagy-related genes including Beclin-1, ATG3, and -5. Moreover, we observed alterations in the fatty acyl composition of cardiolipins and phosphatidic acids. Given the previously reported roles of these phospholipids and Nix in mitochondrial dynamics, we investigated aberrant mitochondrial fission and turn-over as a mechanism of myocyte insulin resistance. In a series of gain-of-function and loss-of-function experiments in rodent and human myotubes, we demonstrate that Nix accumulation triggers mitochondrial depolarization, fragmentation, calcium-dependent activation of DRP1, and mitophagy. In addition, Nix-induced mitochondrial fission leads to myotube insulin resistance through activation of mTOR-p70S6 kinase inhibition of IRS1, which is contingent on phosphatidic acids and Rheb. Finally, we demonstrate that Nix-induced mitophagy and insulin resistance can be reversed by direct phosphorylation of Nix by PKA, leading to the translocation of Nix from the mitochondria and sarcoplasmic reticulum to the cytosol. These findings provide insight into the role of Nix-induced mitophagy and myocyte insulin resistance during an overfed state when overall autophagy-related gene expression is reduced. Furthermore, our data suggests a mechanism by which exercise or pharmacological activation of PKA may overcome myocyte insulin resistance.

**Graphical Abstract:** 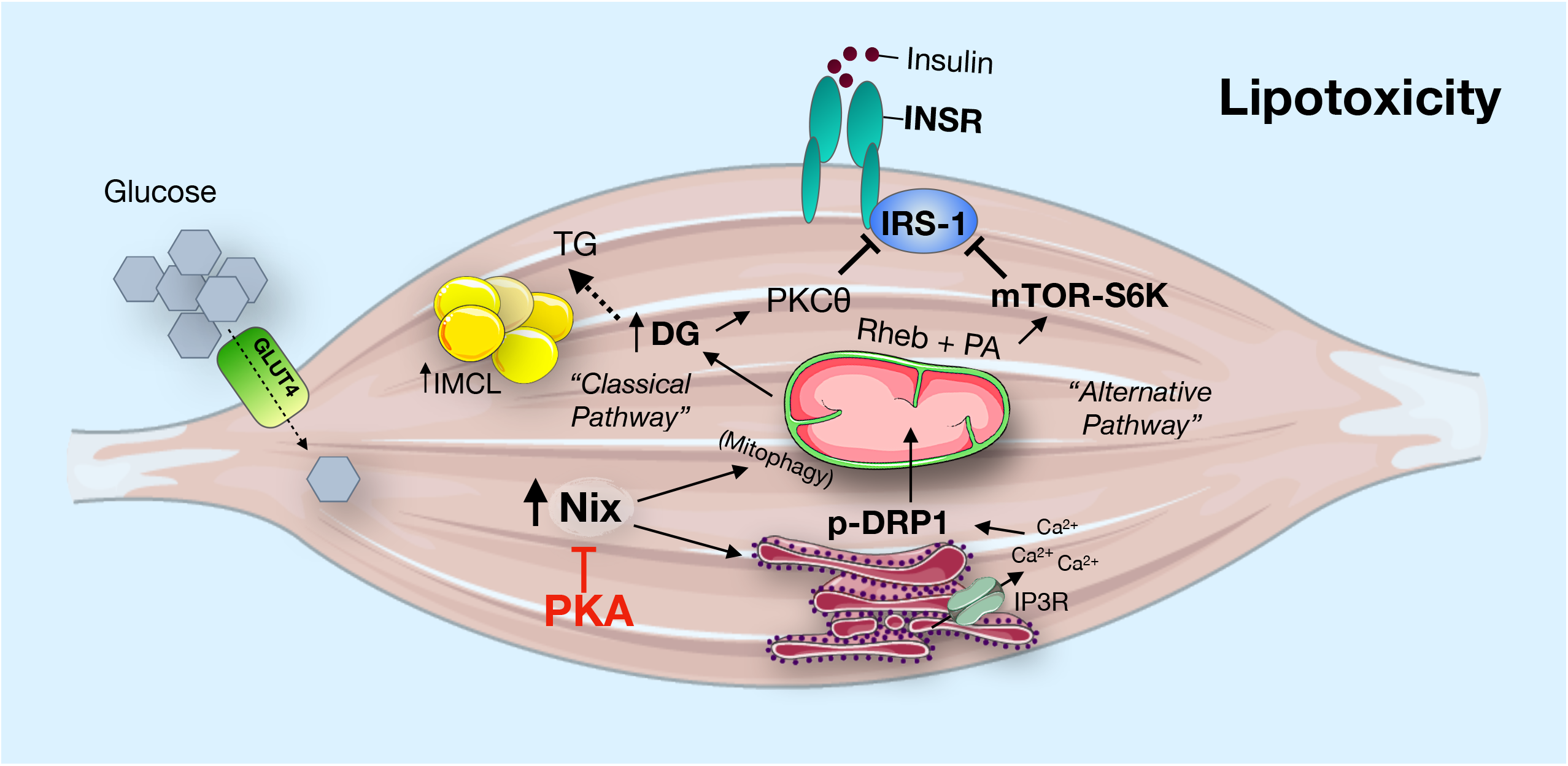

## Introduction

Skeletal muscle insulin resistance is one of the earliest detectable perturbations in the natural progression of type 2 diabetes, as muscle insulin resistance often proceeds and may contribute to hepatic steatosis, as well as adipocyte and beta cell dysfunction (reviewed by^1–3^). Although the cellular mechanisms responsible for muscle insulin resistance have been historically challenging to uncover, and are likely unique from the mechanisms responsible for insulin resistance in other tissues, ectopic lipid accumulation and mitochondrial dysfunction appear to be key events leading to muscle insulin resistance in humans and in rodent models of obesity and type 2 diabetes^1–3^. However, the exact nature of this mitochondrial defect, and how mitochondrial dysfunction impacts insulin signaling and glucose uptake in muscle, remain poorly understood.

Originally proposed by Randle and colleagues, the glucose fatty-acid cycle is a metabolic process involving the competition of glucose and fatty acids for metabolism^4^. More recently, this idea has been expanded to suggest that peripheral insulin resistance involves metabolic inflexibility, where mitochondrial metabolism fails to switch from predominate lipid oxidation to carbohydrate metabolism when faced with the need for glucose disposal^5^. Muscle tissue is likely central to this concept as approximately 80% of postprandial glucose uptake occurs in muscle^6^. Mechanistically, metabolic inflexibility has been attributed to fatty acid-induced mitochondrial overload with depletion of tricarboxylic acid cycle intermediates^7^. An alternative, yet not entirely exclusive hypothesis, involves signaling pathways originating from the toxic accumulation of lipid within a myocyte. Lipotoxicity is a form of cellular stress caused by the accumulation of lipid intermediates resulting in insulin resistance^1,3,8^. Lipid infusion studies in humans, and high-fat feeding in rodents have demonstrated that muscle tissue rapidly accumulates diacylglycerols, ceramides, and triglycerides in response to these lipid exposures^(1,3^. Accumulation of ceramides is thought to active inflammatory JNK signaling, while accumulating diacylglycerols activate novel PKC signaling, including PKCδ and PKCθ^9–13^. Importantly, PKCθ has been shown in both human and rodent studies to phosphorylate and inhibit the insulin receptor substrate-1 (IRS1) to prevent insulin-mediated AKT2 activation and GLUT4-dependent glucose uptake, where a key residue involved in human IRS1 inhibition is Ser-1101^14^. This pathway may protect myocytes from metabolic stress, by preventing further influx of glucose and/or lipids^15^. Intriguingly, acute exercise in humans, which enhances mitochondrial function, reverses muscle insulin resistance concurrent with reduced diacylglycerols and ceramides, and increased triglyceride storage, suggesting diacylglycerols and ceramides are critical lipid intermediates involved in muscle lipotoxicity^9^.

Autophagy is a lysosomal degradation pathway that functions in organelle and protein quality control. During cellular stress, increased levels of autophagy permit cells to adapt to changing nutritional and energy demands through catabolism^16–20^. Although originally described as a cellular response to starvation, autophagy also protects against insulin resistance in fed animals^21^. Acute exercise induces autophagy in skeletal muscle, while transgenic mice harboring a Bcl2 mutant preventing autophagy induction display decreased exercise tolerance and altered glucose metabolism during acute exercise^21^. Furthermore, exercise fails to protect these mice from high fat diet-induced insulin resistance^21^. More recent evidence suggests that autophagy can be selectively targeted to specific cellular structures, such as mitochondria, and mitochondrial proteins that contain an LC3 interacting region (LIR) can serve as selective mitochondrial autophagy receptors (ie. Mitophagy receptors)^(22,23^. A recent study has demonstrated that a skeletal muscle restricted deletion of a hypoxia-inducible mitophagy receptor, called Fundc1, blunts the mitophagy response and protects against high fat feeding-induced insulin resistance at the expense of exercise tolerance, suggesting that excessive mitophagy may be a contributing factor in muscle insulin resistance^24^. However, Fundc1 is hypoxia-inducible and has not been shown to be activated by lipotoxicity; thus, other mitophagy receptors may serve to trigger selective autophagy of dysfunctional mitochondria in a fed lipotoxic state when generalized macro-autophagy is inhibited^(25,26^.

Mitochondrial quality control also involves dynamic fission and fusion events that serve to compartmentalize dysfunctional or depolarized mitochondrial fragments that can be targeted for autophagic degradation^(23,27^. Mitochondrial fission and fusion are regulated by a family of GTPases, where DRP1 initiates mitochondrial fission, and mitofusin-1, -2, and Opa1 regulate fusion^(23,27^. Moreover, lipotoxicity-induced muscle insulin resistance has been associated with excessive mitochondrial fission and activation of DRP1^28^. Interestingly, phospholipids such as phosphatidic acid and cardiolipin, regulate mitochondrial fission and fusion, where cardiolipin facilitates fusion through and interaction with Opa1, and mitochondrial phosphatidic acids interact with DRP1^29–31^. In muscle, phosphatidic acids also regulate mTOR signaling during times of growth^32^. mTOR signaling potently inhibits macro-autophagy in a fed state, and is involved in a negative-feedback pathway that limits glucose uptake through p70S6 kinase (p70S6K)-mediated phosphorylation of IRS1^(33,34^. However, it remains to be determined how these interconnected signaling pathways contribute to muscle insulin resistance.

Previously, we described a lipotoxicity-activated signaling pathway leading to increased expression of the mitophagy receptor Nix^35^. This conserved pathway responds to elevated diacylglycerols by activating PKCδ which inhibits the expression of microRNA-133a, a negative regulator of Nix expression. Furthermore, we established that Nix expression is elevated in muscle tissue of rodents fed a high fat diet (HF)^35^.

In order to further explore the role of Nix and mitochondrial dysfunction associated with muscle insulin resistance, we undertook an unbiased gene expression array and a shot-gun metabolomics screen in soleus muscle from rodents on a low fat (LF) or HF diet. We observed a decrease in the most abundant tetralinoleoyl-cardiolipin species and an increase in phosphatidic acids, concurrent with increased Nix expression, suggesting altered mitochondrial dynamics and mitophagy contribute to muscle insulin resistance. Using two myocyte cell lines, and human myotubes differentiated from induced pluripotent stem cells (iPSCs), we mechanistically determined that Nix accumulation triggers calcium-dependent activation of DRP1 and mitophagy. In addition, Nix-induced mitochondrial fission leads to impaired insulin signaling in myotubes through mTOR-dependent inhibition of IRS1. Finally, we demonstrate that Nix-induced mitophagy and impaired insulin signaling can be reversed by direct phosphorylation of Nix by PKA, leading to the translocation of Nix from the mitochondria and sarcoplasmic reticulum to the cytosol. These findings are consistent with a model whereby Nix responds to myocyte lipotoxicity in order to clear damaged mitochondria through receptor-mediated mitophagy, and protect the myocyte against nutrient storage stress by activating mTOR-dependent desensitization of insulin signaling.

## Results

Numerous lipid species were elevated in soleus muscle with HF feeding, including diacylglycerols, ceramides, and triglycerides (Table 1 and Supplemental tables). Interestingly, we also observed alterations in the composition of cardiolipin. Cardiolipin is normally found in the inner mitochondrial membrane and uniquely contains four acyl chains, where the tetra-linoleoyl (18:2) cardiolipin is the most abundant species. Interestingly, we observed a reduction in this species of cardiolipin with HF feeding (Table 1). We also observed an increase in most phosphatidic acid species, where the largest increases were observed in PA(20:0/22:6) and PA(19:0/20:0)(Table 1). To further explore the relationship between lipotoxicity and skeletal muscle autophagy, we performed a PCR-based array on soleus muscle from rats fed a LF or HF diet. We observed a modest reduction in several autophagy-related genes including Beclin-1, ATG3, -5, -12 and Atp6v1g2 in HF fed animals (Table 1 and Supplemental tables). These observations suggested that global autophagy-related gene expression is reduced in muscle during an over-fed state. Interestingly, we also observed a characteristic reduction in PGC-1a, and a corresponding reduction in mitochondrial enzymes, such as citrate synthase (CS) and malate dehydrogenase (MDH1). However, other mitochondrial marker genes, such as translocases of the outer and inner mitochondrial membrane (TOM and TIM complexes) were relatively unchanged, suggesting an alteration in the mitochondrial phenotype rather than a simple reduction in mitochondrial content. Perhaps surprisingly, we observed reductions in genes typically associated with the red (ie. slow twitch) muscle fibre-type, such as myosin heaving chains-1 and -2 (Myh1, Myh2), myoglobin (Mb), and troponin- C1 and -T1 (Tnnc1, Tnnt1). In addition, we observed an increase in myostatin (Mstn), and an increase in myogenesis transcription factors such as Myf5 and MyoD1.

**Table 1.** Representative metabolites and mRNAs altered by HF feeding.

| **Metabolite** | **Fold Change** |
| --- | --- |
| **Triacylglyceride** |  |
| TG(22:3/22:6/22:6) | 474 |
| TG(21:0/22:0/22:3) | 225 |
| TG(20:0/22:0/22:6) | 252 |
| **Diacylglyceride** |  |
| DG(22:3/22:6) | 315 |
| DG(22:2/24:1) | 685 |
| DG(20:5/22:4) | 380753 |
| DG(20:4/22:6) | 6177 |
| DG(20:1/18:0) | 389 |
| DG(20:0/22:1) | 209 |
| DG(18:4/22:6) | 500 |
| DG(16:0/18:0) | 8408 |
| **Ceramides** |  |
| Ceramide (d18:1/26:0) | 205 |
| Cer(t18:0/16:0) | 5830 |
| Cer(d18:2/20:1) | 298 |
| Cer(d18:2/14:0) | 283 |
| Cer(d16:2/20:1) | 122962 |
| **Cardiolipins (CL)** |  |
| CL(18:2/18:2/18:2/18:2) | 0.003 |
| **Phosphatidic Acids (PA)** |  |
| PA(20:0/22:6) | 19843 |
| PA(20:0/18:2) | 510 |
| PA(20:3/21:0) | 518 |
| PA(19:0/20:0) | 12389 |
| **Genes** | **Fold Change** |
| Atg12 | 0.82 |
| Atg3 | 0.86 |
| Atg5 | 0.73 |
| Atp6v1g2 | 0.39 |
| Becn1 | 0.84 |
| Cs | 0.80 |
| Mb | 0.48 |
| Myh1 | 0.38 |
| Ppargc1a | 0.65 |

Given our previous findings that Nix is induced in this rodent model^35^, and that phospholipids have been implicated in both mitochondrial dynamics and mTOR signaling, we investigated the role of Nix in regulating aspects of mitochondrial quality control following exposure to lipotoxicity. To begin, we expressed Nix in C2C12 myoblasts and monitored mitochondrial morphology using a mitochondrial-targeted Emerald fluorophore (mito-emerald). In control cells we observed predominately elongated mitochondria; however, in cells expressing Nix we observed a decrease in elongated mitochondria and increases in mitochondria with an intermediate or ‘fissioned’ morphology and increases in overt mitochondrial fragmentation with a more pronounced perinuclear distribution (Figure 1A, -B). In addition, expression of Nix in both C2C12 and L6 myoblasts resulted in a significant decrease in mitochondrial membrane potential, determined by TMRM staining (Figure 1C, -D). Previously, we demonstrated that Nix-induced mitochondrial depolarization in cardiomyocytes was due to ER/SR-dependent calcium release and subsequent mitochondrial permeability transition^36^. Thus, we examined both steady-state ER/SR and mitochondrial calcium content in cells expressing Nix using organelle-targeted calcium biosensors called GECOs^37–39^. We observed that Nix expression reduced steady-state ER/SR calcium and increased steady-state mitochondrial calcium (Figure 1E, -F). Furthermore, treatment of C2C12 cells with an IP3-receptor blocker (2APB) prevented Nix-induced mitochondrial depolarization (Supplemental Figure 1A). Previous work has identified that the mitochondrial fission protein DRP1 is activated by calcineurin, a calcium-calmodulin dependent phosphatase^40^. Thus, we determined if Nix-induced ER/SR calcium release could result in DRP1 dephosphorylation at a known calcineurin site (Mouse Ser643; Human Ser637). In C2C12 cells expressing Nix, we observed a reduction in Ser643 phosphorylation, compared to control cells (Figure 1G). As mitochondrial fission and loss of membrane potential are important precursor events leading to mitochondrial autophagy, we evaluated mitophagy using a mitochondrial matrix-targeted pH biosensor called mito-pHred^41^. Nix expression in L6 and C2C12 myoblasts and myotubes increased mito-pHred fluorescence (Figure 1H-J). As positive and negative controls, we expressed Parkin and mito-pHred and observed increased fluorescence, while expression of a dominate-negative ATG5 prevented Nix-induced mito-pHred activation (Supplemental Figure 1A-B). Moreover, Nix-induced mito-pHred activation was prevented by a dominate-negative DRP1, the mitochondrial fission inhibitor mdivi-1, and the lysosomal inhibitor Bafalomycin-A1 (Figure 1K-L).

**Figure 1.**
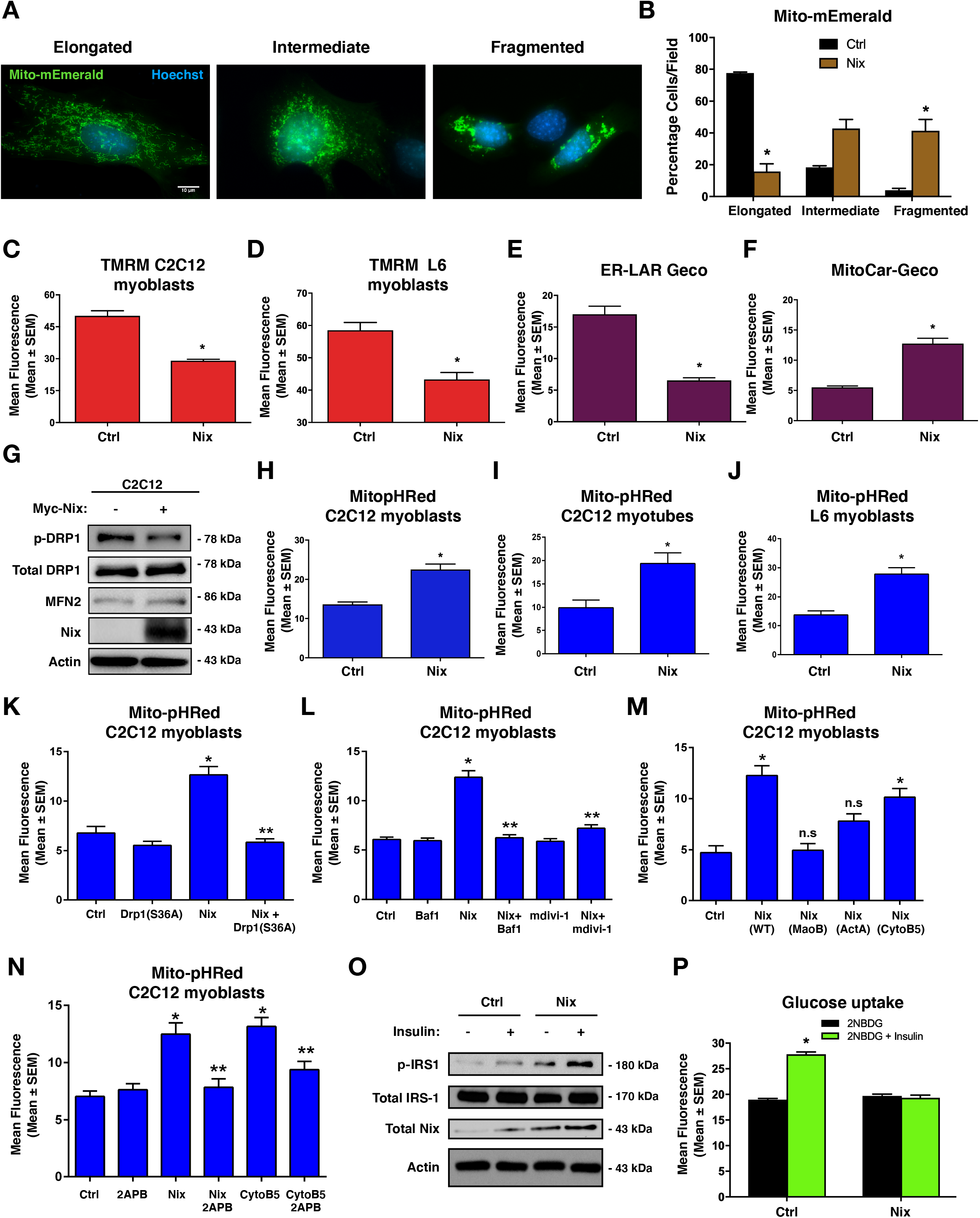
Nix regulates mitochondrial dynamics and mitophagy. A-B) C2C12 myoblast cells were transfected with mito-mEmerald to assess mitochondrial morphology (left). Quantification of C2C12 myoblast cells transfected Nix or empty vector control (right). C-D) Quantification of C2C12 myoblast cells (c) L6 myoblasts (d) transfected with Nix or an empty vector control; stained with TMRM. E) Quantification of C2C12 myoblasts transfected with ER-LAR-GECO, Nix, or an empty vector control. F) Quantification of C2C12 myoblasts transfected with Mito-CAR-GECO, Nix, or an empty vector control. G) C2C12 cells were transfected with Nix or empty vector. Protein extracts were analyzed as indicated. H-J) Quantification of (h) C2C12 myoblasts, (i) C2C12 myotubes, and (j) L6 myoblasts cells transfected with Nix, mito-pHRed, or an empty vector control. K) C2C12 cells were transfected with mito-pHRed, Nix, a dominant negative Drp1 (S36A), or an empty vector control and quantified. L) C2C12 cells were transfected with mito-pHRed, Nix, or an empty vector control. Cells were treated with Bafilomycin (6nM, 3h) or with the mitochondrial fission inhibitor (mdivi-1, 20μM, 1h). M) Quantification of C2C12 myoblast cells transfected with mito-pHRed, Nix, mitochondrial targeted Nix fusion constructs Nix-MaoB or Nix-ActA, the ER/SR targeted Nix constuct Nix-CytoB5, or an empty vector control. N) C2C12 myoblast cells were transfected with mito-pHRed, Nix, Nix-CytoB5, or an empty vector control. Cells were treated with 2-aminoethoxydiphenyl borate (2APB 10μM, 1h), or DMSO as control vehicle and quantified. O) L6 myotubes were transfected with Nix or empty vector followed by 15 minutes of insulin stimulation (10nM). Protein extracts were analyzed as indicated. P) L6 myotubes were transfected with Nix or an empty vector control. Insulin stimulated glucose uptake (10nM) was determined by 2NBDG fluorescence and quantified. Data are represented as mean ± S.E.M. *P < 0.05 compared with control, while **P < 0.05 compared with treatment, determined by 1-way or 2-way ANOVA.

To further delineate the role of Nix as a mitophagy receptor at the mitochondria and as a regulator of calcium release at the ER/SR in our models, we utilized organelle-targeted Nix constructs, as described previously^36^. Interestingly, mitochondrial targeted Nix constructs, using either the monoamine oxidase-B (Nix-MaoB) or the *Listeria* ActA (Nix-ActA) targeting domains, had little to no effect on mito-pHred activation; whereas, ER/SR targeted Nix using the cytochrome-B5 (Nix-CytoB5) targeting domain had a similar effect to wild-type Nix (Figure 1M). In addition, we observed that both Nix and Nix-CytoB5-induced mito-pHred activation was prevented with the IP3-receptor blocker 2APB. Finally, we observed that Nix expression led to enhanced phosphorylation of IRS1 using the phospho-specific Ser-1101 antibody (Figure 1O), and prevented insulin-stimulated glucose uptake, determined by 2NBDG fluorescence (Figure 1P). Collectively, our findings suggest that in addition to its role as a mitophagy receptor, Nix regulates mitochondrial fission through its effects on ER/SR calcium, and is also a regulator of insulin sensitivity.

Next, we confirmed that Nix expression is elevated in soleus muscle of rats fed a HF diet compared to those fed a LF diet, concurrent with decreased phosphorylation of DRP1 (Figure 2A). Interestingly, other markers of mitophagy, such as Bnip3, Bcl2l13, Fundc1, and Rheb, were unchanged or modestly decreased in HF fed rodents, while Fkbp8 and Parkin were modestly increased (Supplemental Figure 1D). We also evaluated if Parkin activation could be downstream of Nix induction, but Nix expression failed to promote the mitochondrial localization of Parkin-YFP (Supplemental Figure 1E). In culture, we used palmitate treatment to induce lipotoxicity and performed a dose-response in C2C12 cells. Nix expression was increased by 0.2 mM palmitate and plateaued at higher concentrations (Figure 2B). Furthermore, palmitate treatment increased mito-pHred fluorescence, which was inhibited by knock-down of Nix (shNix)(Figure 2C). Knockdown of Nix also prevented palmitate-induced ER/SR calcium release and mitochondrial calcium accumulation (Figure 2D-E), which were also blocked by 2APB treatment (Supplemental Figure 1F-G). Furthermore, palmitate treatment decreased DRP1 phosphorylation at the calcineurin regulated residue and led to mitochondrial fission, which were restored by shNix (Figure 2F, Supplemental Figure 1H). In addition, Nix knock-down prevented palmitate-induced mitochondrial depolarization in both C2C12 cells and iPSC-derived human myotubes (Figure 2G-H), and restored insulin-stimulated glucose uptake following palmitate exposure (Figure 2I). Using a diacylglycerol biosensor called DAGR^42^, we observed that palmitate treatment increased FRET emission, which was reversed by Nix knock-down (Figure 2J), suggesting a connection between mitochondrial function and diacylglycerol accumulation. To test the specificity of shNix, we treated C2C12 myoblasts with palmitate and observed an increase in mito-pHred fluorescence, which was prevented when cells expressed shNix, but was restored by coexpression of a Myc-tagged Nix (Myc-Nix)(Figure 2K). Finally, we observed that palmitate-induced mitophagy was prevented by 2APB and with Etomoxir, an inhibitor of mitochondrial fatty acid uptake through carnitine palmitoyltransferase-1 (Figure 2L), consistent with the notion that fatty acid-induced mitochondrial overload may be important trigger for mitophagy.

**Figure 2.**
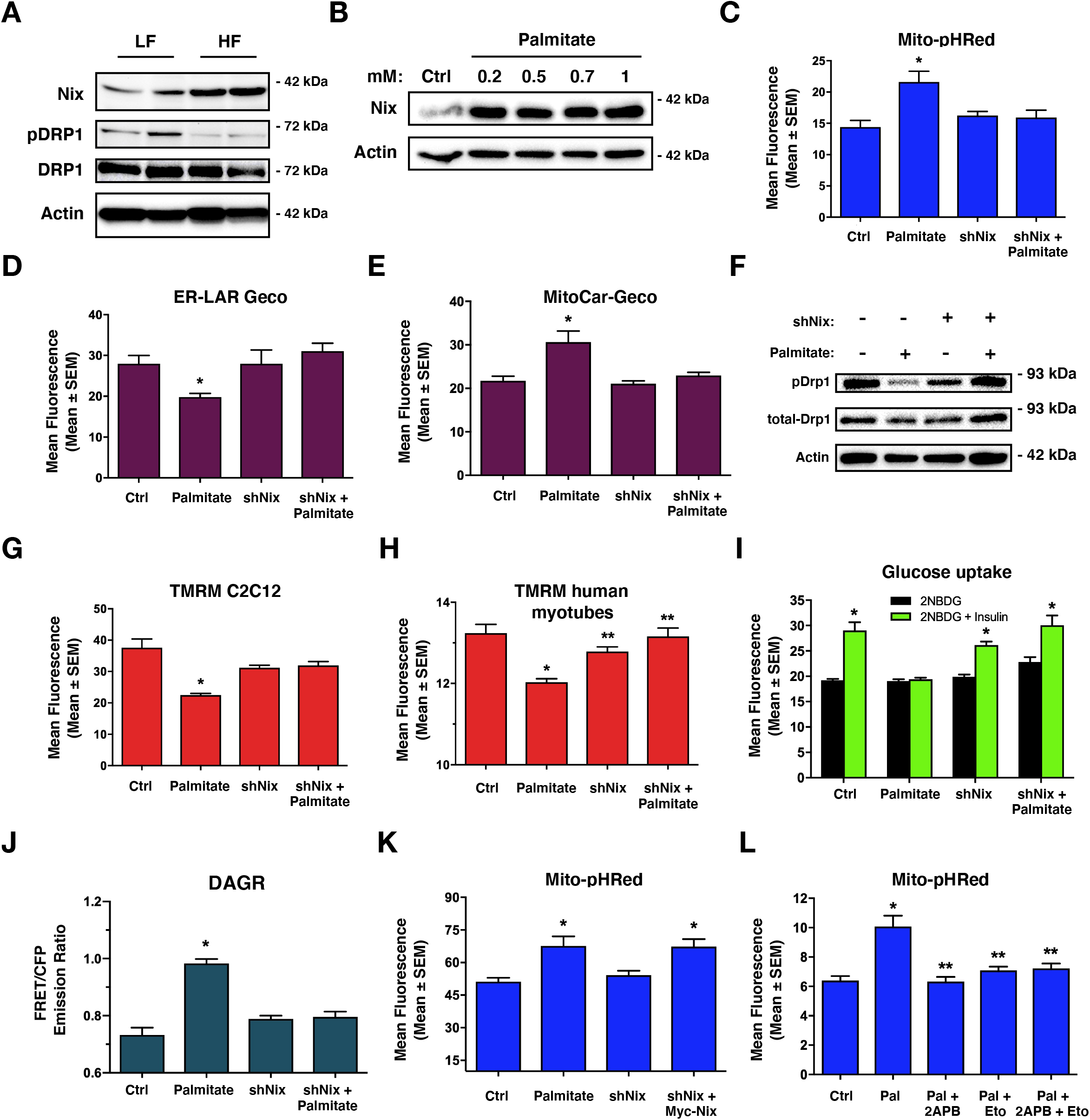
Knockdown of Nix improves mitochondrial function and insulin sensitivity. A) Western blot analysis of rat soleus muscle exposed to high fat (HF) or low fat (LF) diet for 12-weeks. B) C2C12 myoblast cells were treated overnight with increasing doses of palmitate conjugated to 2% albumin in low glucose media. Protein extracts were analyzed as indicated. C) C2C12 myoblast cells were transfected with mito-pHRed, shNix, or a scrambled control shRNA. Cells were treated overnight with palmitate (200μM). D) Quantification of C2C12 myoblasts transfected with ER-LAR-GECO and treated as in (c). E) Quantification of C2C12 myoblast cells transfected with Mito-CAR-GECO and treated as in (c). F) C2C12 myoblast cells were transfected with shNix or a scrambled control shRNA, followed by palmitate treatment as described in (c). Protein extracts were analyzed as indicated. G) C2C12 myoblast and H) human myotube cells were transfected with shNix or a control shRNA. Cells were treated overnight with palmitate as in (c), stained for TMRM and quantified. I) L6 myotubes were transfected with shNix or a control shRNA. Cells were treated overnight with palmitate as in (c). Insulin stimulated glucose uptake (10 nM) was determined by 2NBDG fluorescence and quantified. J) C2C12 myoblast cells as transfected as in (h), along with DAGR, a diacylglycerol biosensor. Cells were analyzed by using the emission ratio of YFP to CFP (FRET Ratio). K) Quantification of C2C12 myoblast cells transfected as in (h), along with Myc-Nix and treated as in (c). L) C2C12 cells were transfected with mito-pHRed. Cells were pre-treated overnight with etomoxir (100μM) and palmitate, as described in (c), followed by co-treatment with 2APB (10μM, 2h) or DMSO as control vehicle and quantified. Data are represented as mean ± S.E.M. *P < 0.05 compared with control, while **P < 0.05 compared with treatment, determined by 1-way or 2-way ANOVA.

Next, we explored whether Nix function was directly regulated by cellular signaling pathways to modulate mitophagy and insulin sensitivity. *In silico* analysis of the Nix amino acid sequence identified a conserved PKA consensus motif within the carboxy-terminus of the transmembrane domain (Figure 3A-B), which is Serine-212 in human Nix. We engineered synthetic peptides spanning the Nix transmembrane domain and subjected them to *in vitro* kinase reaction with the catalytic subunit of PKA and analyzed peptides by ion trap mass spectrometry. In the absence of kinase, single ion monitoring (SIM) scans displayed a predominant peak at m/z of 857.28 (z=2+)(Figure 3C). Following kinase incubation, this peak shifted by m/z of 40 (897.78, z=2+), corresponding to the addition of a phosphate (PO_3_) to the peptide (Mass = 80.00 Da)(Figure 3D). However, when Serine-212 was mutated to a neutral alanine (S212A) this m/z shift was eliminated (Figure 3E). Next, we analyzed the MS^2^ spectra produced by collision-induced dissociation (CID) of the precursor ion with m/z = 897.78 (z=2+). CID of the phospho-peptide yielded a product-ion with m/z of 848.67 (delta = 49.11), consistent with the neutral loss of phosphorylation (98/2 = 49; Figure 3F). We also evaluated if our Nix peptide could be phosphorylated at more than one site; however, we did not detect an m/z shift equivalent to 160 Da (Figure 3G). We modeled the Nix structure using the Phyre^2^ engine (Figure 3H). Using known 3D motifs, Phyre^2^ predicted that Serine-212 was exposed within the transmembrane domain, and likely well-positioned for kinase recognition.

**Figure 3.**
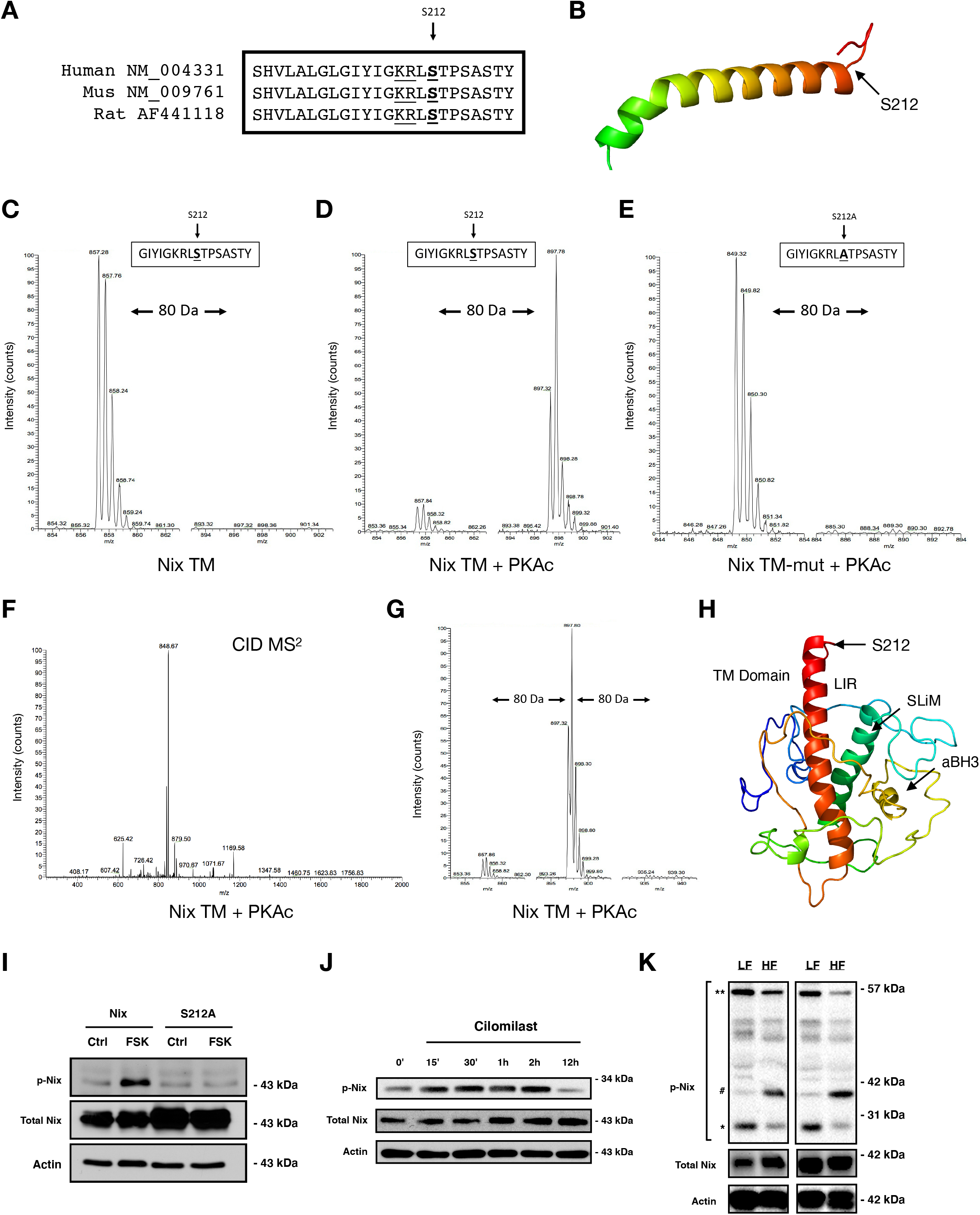
PKA phosphorylates Nix at serine-212. A) Amino acid sequence alignment for human, mouse and rat Nix. B) Schematic Nix transmembrane domain and the location of Serine-212. C) SIM scan of the wild-type peptide spanning the PKA site of Nix. The unphosphorylated peptide has 857.28 m/z (z=2+). D) Putative phosphorylation showing an increased m/z of 40 that corresponds to PO_3_ (M=80.00 Da). E) SIM scan of mutated peptide (S212A) incubated with kinase, as in (d). F) MS^2^ spectra following collision-induced dissociation of the precursor ion in (D, m/z=897.78). A neutral loss of 49 m/z confirms phosphorylation. G) SIM scanning showing Nix peptide is not phosphorylated at two residues within the peptide. H) Nix structure modeled using Phyre2 engine. I) 3T3 cells were transfected with wild type Nix or a Nix mutant where Serine-212 is converted to alanine (S212A) and treated with 10μM forskolin for 2 hours. Protein extracts were immunoblotted, as indicated. J) C2C12 myoblast cells were treated with 10 μM cilomilast or vehicle for multiple time points. Protein extracts were immunoblotted, as indicated. K) Phospho-Nix is decreased as total Nix increases in soleus muscle of rodents treated with high fat (HF) or low fat (LF) diet for 12-weeks. *predicted molecular weight, **predicted dimer weight, ^#^aligned with total Nix.

Next, we used a custom antibody designed to specifically detect phosphorylated Serine-212 of Nix (pNix)(Abgent). Following treatment of C2C12 cells with forskolin (FSK) or a cAMP analog, we detected a band that aligned with transfected Myc-tagged Nix (Figure 3I and Supplemental Figure 2A). However, when Serine-212 was mutated to a neutral alanine, this band was lost (Figure 3I). While investigating the endogenous expression of pNix, we observed a notable cell-type specific pattern by western blot. Nix has a predicted molecular weight of 26 kDa, but has been observed to migrate on SDS-PAGE as a 40 kDa monomer, and an 80 kDa dimer^35,43,44^. In both C2C12 and L6 myoblasts, the dominant pNix band migrated close to the predicted weight of Nix (Supplemental Figure 2B-D). This band was also sensitive to H89 treatment and was substantially reduced in cells expressing an shRNA targeting Nix (Supplemental Figures 2C-D). However, in the human rhabdomyosarcomal cell lines (RH30 and A204), the pNix antibody detected multiple bands spanning 40-80 kDa, while in human iPSC-derived myoblasts the dominant band migrated at the predicted dimer weight (Supplemental Figure 2B). These observations suggest that Nix is likely exposed to other post-translational modifications, and that these modifications occur in a cell-type or species-specific manner. Next, we performed a series of time-courses experiments with pharmacological agents known to activate PKA signaling in muscle. We used the adrenergic agonist clenbuterol and the phosphodiesterase-4 inhibitor cilomilast, and confirmed they activated PKA in C2C12 myoblasts (Supplemental Figure 2F-G). Both of these agents increased pNix expression peaking at 2-hours and returning to control levels by 12-18 hours (Figure 3J and Supplemental Figure 2E). In addition, expression of PKA and treatment of C2C12 cells with clenbuterol prevented Nix-induced mitochondrial depolarization (Supplemental Figure 3). Lastly, we probed soleus muscle extracts from rodents fed a LF or HF diet. In rodent muscle we observed three dominant pNix bands; a predicted 26 kDa, a 52 kDa dimer band, and a 40 kDa band that aligned with total Nix. The 40 kDa band is likely more detectable in muscle tissue as total Nix expression is much greater in muscle tissue compared to C2C12 and L6 myoblasts. In addition, we observed that the 26 kDa band and the dimer band were reduced with HF feeding (Figure 3K), while the 40 kDa band more closely followed total Nix expression. Based on the positioning of the phospho-residue within the transmembrane domain of Nix, we hypothesized that Ser-212 is an inhibitory phosphorylation site.

Next, we performed a series of experiments to evaluate if clenbuterol and cilomilast could inhibit palmitate-induced mitochondrial defects. In both C2C12 and L6 myotubes, clenbuterol and cilomilast reversed palmitate-induced mitochondrial depolarization (Fig 4A-B). We also evaluated human iPSC-derived myotubes and observed similar rescue of TMRM and mito-Sox staining with clenbuterol and cilomilast treatment (Fig 4C-E). In addition, we confirmed that clenbuterol and cilomilast increase pNix expression in human myotubes (Fig 4F). Clenbuterol and cilomilast also prevented palmitate-induced alterations in mitochondrial morphology, mitophagy, insulin signaling, and insulin-stimulated glucose uptake (Fig 4G-J). Using TMRM as an indicator of Nix activity, we observed that clenbuterol reversed Nix-induced mitochondrial depolarization, but not when Ser-212 was mutated to alanine (S212A; Fig 4K). In addition, we generated a Ser-212 phospho-mimetic mutation (S212D) and compared the effects of this construct to wild-type Nix and the Nix-S212A mutant. When transfected into C2C12 cells, wild-type Nix and Nix-S212A induced mitophagy, increased mitochondrial calcium, reduced ER/SR calcium, and inhibited insulin-stimulated glucose uptake (Fig 4L-O). However, the Nix-S212D mutant did not significantly impact these end-points. Collectively, these findings suggest that clenbuterol and cilomilast treatment leads to PKA-induced phosphorylation of Nix at Ser-212, and this phosphorylation site is inhibitory and can modulate Nix-induced mitophagy, calcium homeostasis, and insulin sensitivity.

**Figure 4.**
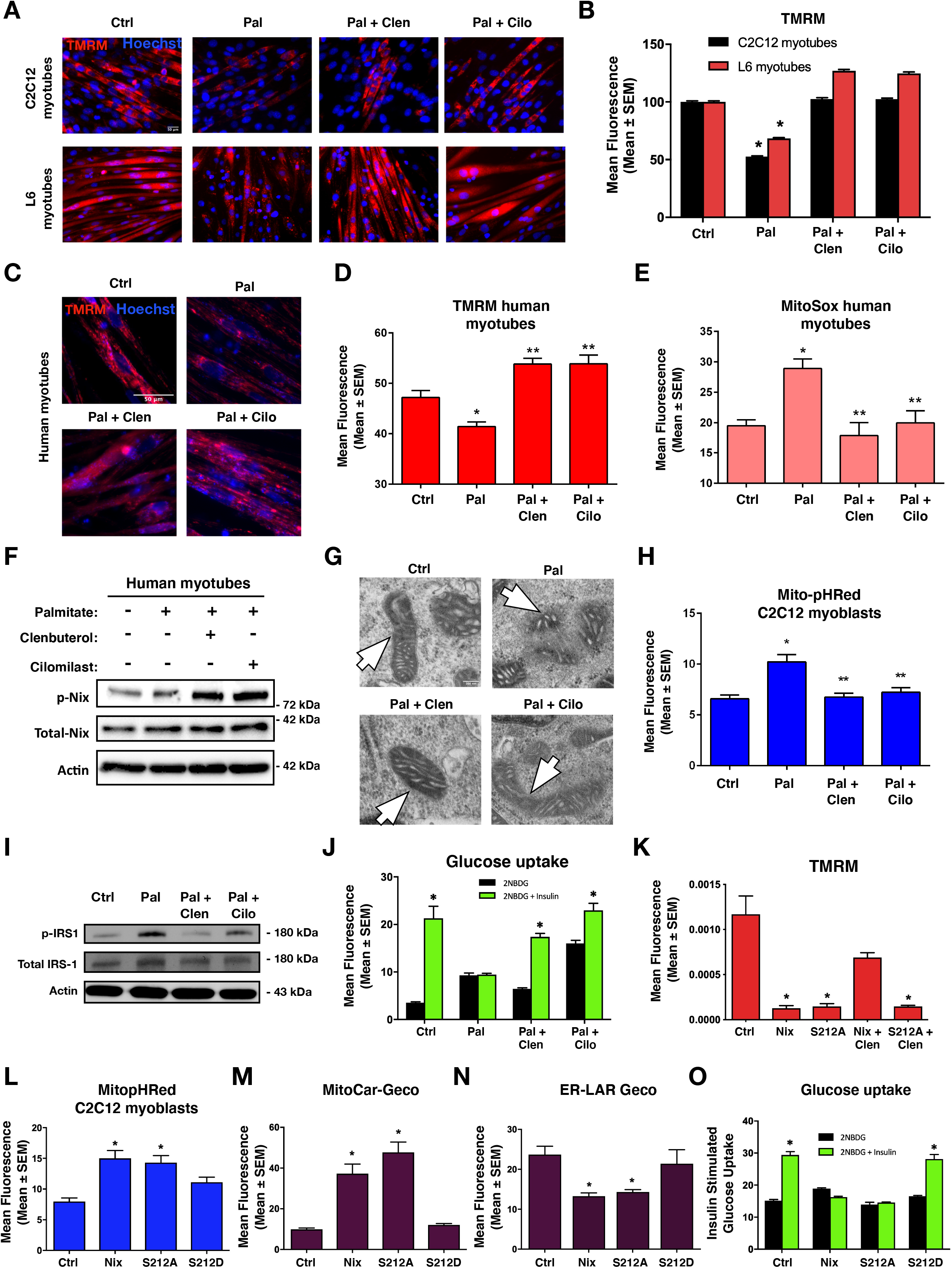
Clenbuterol and cilomilast inhibit palmitate-induced mitochondrial defects. A) 5-day differentiated C2C12 myotubes and L6 myotubes were treated overnight with palmitate (200μM) conjugated to 2% albumin in low glucose media. Control cells were treated with 2% albumin alone. Myotubes were treated with clenbuterol (500nM, 2h), cilomilast (10μM, 2h) or DMSO control vehicle and stained with TMRM (red) and Hoechst (blue) and imaged by standard fluorescent microscopy. B) Quantification of (a). C) 7-day differentiated human myotubes were treated and stained as in (a). D) Quantification of (c). E) 7-day differentiated human myotubes were stained with Mitosox and quantified. F) 7-day differentiated human myotubes were treated as in (a). Protein extracts were immunoblotted, as indicated. G) C2C12 myoblast cells were treated as in (a) and imaged via electron microscopy. Mitochondria are indicated by arrows. H) C2C12 myoblasts were transfected with mito-pHRed and treated as in (a). I) L6 myotube were treated as in (a) and protein extracts were immunoblotted, as indicated. J) Insulin stimulated glucose uptake (10 nM) was determined by 2NBDG fluorescence in treated L6 myotubes and quantified. K) C2C12 myoblast cells were transfected with Nix wild type and Nix-S212A followed by clenbuterol treatment (500nM, 2h). Cells were stained with TMRM and imaged by standard fluorescent microscopy. L-N) C2C12 myoblasts cells were transfected with Nix, Nix-S212A, or Nix-S212D with mito-pHRed (l), mito-Car-Geco (m), and ER-Lar-Geco (n). O) L6 myotubes cells were transfected with Nix, Nix-S212A, or Nix-S212D. Insulin stimulated uptake (10 nM) was determined by 2NBDG fluorescence and quantified. Data are represented as mean ± S.E.M. *P < 0.05 compared with control, while **P < 0.05 compared with treatment, determined by 1-way or 2-way ANOVA.

To determine how PKA phosphorylation of Ser-212 leads to inhibition of Nix-induced mitophagy and impaired insulin signaling, we undertook cell fractionation experiments to determine the cellular localization of pNix. Although the majority of Nix localized to the mitochondria and ER/SR in C2C12 myoblasts, pNix was exclusively localized in the cytosolic fraction (Fig 5A). Moreover, our *In silico* analysis of Nix predicted that Ser-212 lies within a conserved interacting domain of the molecular chaperone family 14-3-3, which identify the sequence RSxpSxP (Nix: KRLpSTP) and are commonly found within PKA and CaMKII phosphomotifs. Thus, we evaluated whether 14-3-3 proteins could translocate Nix from the mitochondria and/or ER/SR upon PKA phosphorylation. First, we performed co-immunoprecipitation of Nix with 14-3-3ß. We chose this 14-3-3 family member as it has been shown to interact with other Bcl-2 proteins^45–47^, and 14-3-3² expression was altered in our gene expression screen of insulin resistant soleus muscle (not shown). We expressed Myc-tagged Nix and HA-tagged 14-3-3² in 293T cells. Shown in Figure 5B, we detected HA-14-3-3² following immunoprecipitation with a Myc antibody. Furthermore, when transfected C2C12 cells were treated with clenbuterol prior to co-immunoprecipitation, the interaction between Nix and 14-3-3² was enhanced (Fig 5C). In a complementary experiment, we expressed HA-14-3-3² with either wild-type Nix, Nix-S21A, or Nix-S212D and subjected extracts to co-immunoprecipitation. We observed that the interaction between Nix and 14-3-3 was enhanced when Ser-212 was mutated to aspartic acid (Nix-S212D)(Fig 5D). Next, we expressed Nix with and without HA-14-3-3² treated with clenbuterol, and subjected extracts to sub-cellular fractionation. We observed that expression of HA-14-3-3² and treatment with clenbuterol reduced the expression of Nix in the mitochondrial and ER/SR fractions, with a corresponding increase in Nix in the cytosolic fraction, and no change in total Nix expression (Fig 5E). Finally, we evaluated if 14-3-3² could counteract the effects of Nix on mitochondria and insulin signaling. Shown in Figure 5F, and -G, co-expression of 14-3-3² with Nix prevented ER/SR calcium release and mitochondrial calcium accumulation. In addition, 14-3-3² blocked Nix-induced mitophagy (Fig 5H), prevented Nix-induced IRS1 phosphorylation (Fig 5I), and restored insulin-stimulated glucose uptake (Fig 5J).

**Figure 5.**
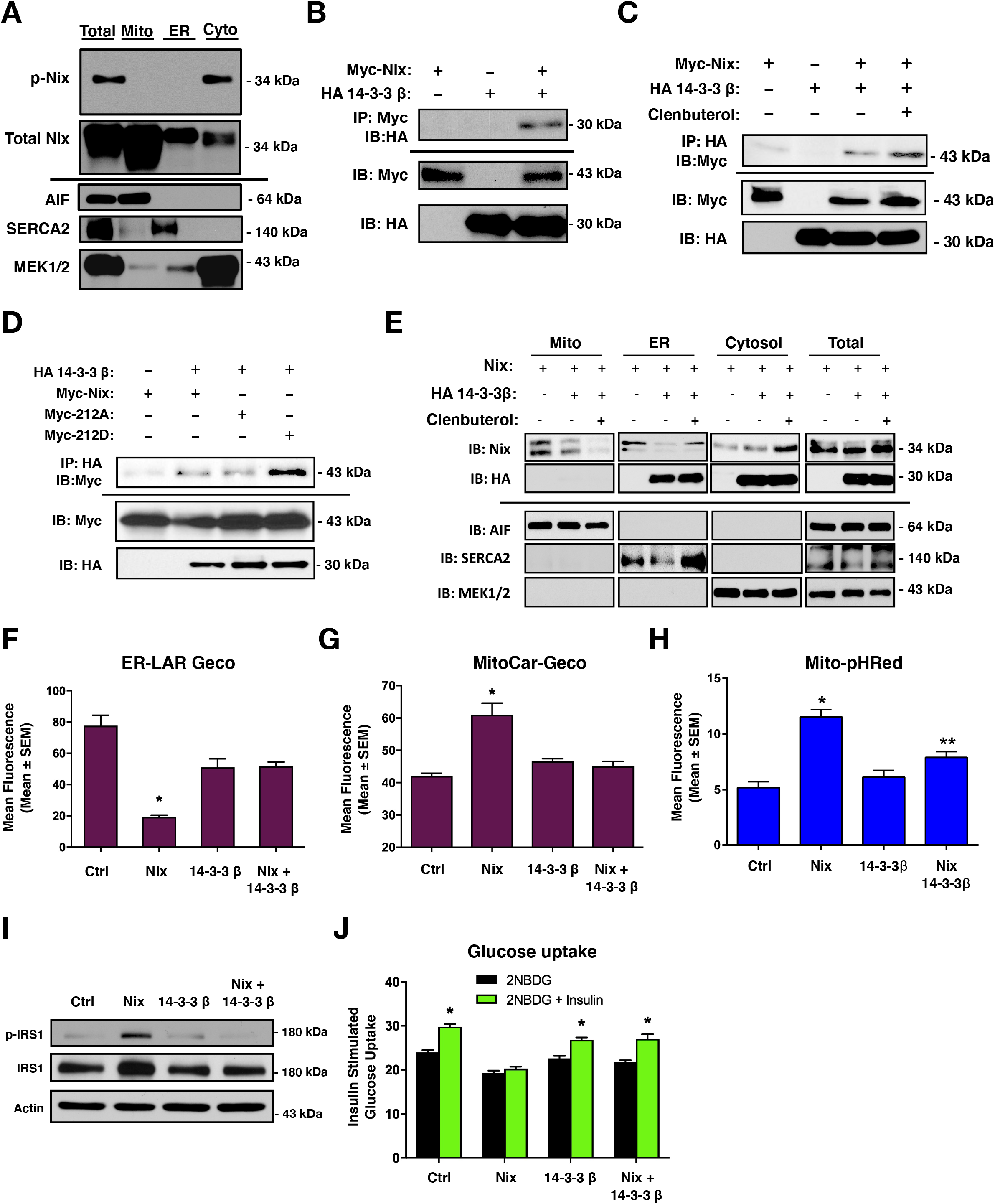
Phospho-Nix interacts with 14-3-3 proteins to determine subcellular location. A) C2C12 myoblasts were subjected to subcellular fractionation. Protein extracts were immunoblotted for phospho-Nix and total Nix, as indicated. B) 293T cells were transfected with Myc-Nix or HA 14-3-3². Extracts were immunoprecipitated (IP) with a Myc antibody and immunoblotted (IB), as indicated. C) C2C12 myoblasts were transfected as in (b) and treated with clenbuterol (500nM, 2h). Extracts were immunoprecipitated (IP) with HA antibody and immunoblotted (IB), as indicated. D) C2C12 myoblasts were transfected with HA 14-3-3², Myc-Nix, Myc-Nix-S212A or Myc-Nix-S212D. Extracts were immunoprecipitated (IP) with HA antibody and immunoblotted (IB), as indicated. E) C2C12 myoblasts were transfected with either Myc-Nix or HA 14-3-3², followed by clenbuterol treatment (500nM, 2h). Extracts were fractioned and protein extracts were immunoblotted, as indicated. F) C2C12 myoblasts cells were transfected with Nix wild type, HA 14-3-3², and ER-Lar-Geco. G) C2C12 myoblasts cells were transfected with Nix wild type, HA 14-3-3², and mitoCar-Geco. H) C2C12 myoblasts cells were transfected with Nix wild type, HA 14-3-3², and mito-pHRed. I-J) L6 myotubes were transfected with Nix wild type, HA 14-3-3². Proteins were immunoblotted as indicated (i) and insulin stimulated glucose uptake (10 nM) was determined by 2NBDG fluorescence and quantified (j). Data are represented as mean ± S.E.M. *P < 0.05 compared with control, while **P < 0.05 compared with treatment, determined by 1-way or 2-way ANOVA.

Previous studies have demonstrated that Ser-1101 is phosphorylated by both novel PKC isoforms and p70S6K to inhibit insulin signaling^13,14,33^. Interestingly, both of these pathways are activated by lipid intermediates, diacylglycerols and phosphatidic acids, respectively, which were elevated in our metabolomics screen of insulin resistant muscle. To investigated how Nix might activate p70S6K signaling, we expressed Nix in C2C12 cells and evaluated activation of p70S6K by phosphorylation at Threonine-389 by phospho-specific antibody. Nix expression increased p70S6K phosphorylation, which was prevented by treatment with the mTORC1 inhibitor rapamycin (Figure 6A). In addition, palmitate treatment increased p70S6K phosphorylation, which was attenuated in cells expressing shNix (Figure 6B). Furthermore, Nix-induced p70S6K phosphorylation was inhibited by clenbuterol and cilomilast treatment (Figure 6C), and prevented by treatment of the mitochondrial fission inhibitor mdivi-1 (Figure 6D), collectively suggesting that Nix-induced mitochondrial fission is necessary and sufficient to activate p70S6K phosphorylation in C2C12 cells.

**Figure 6.**
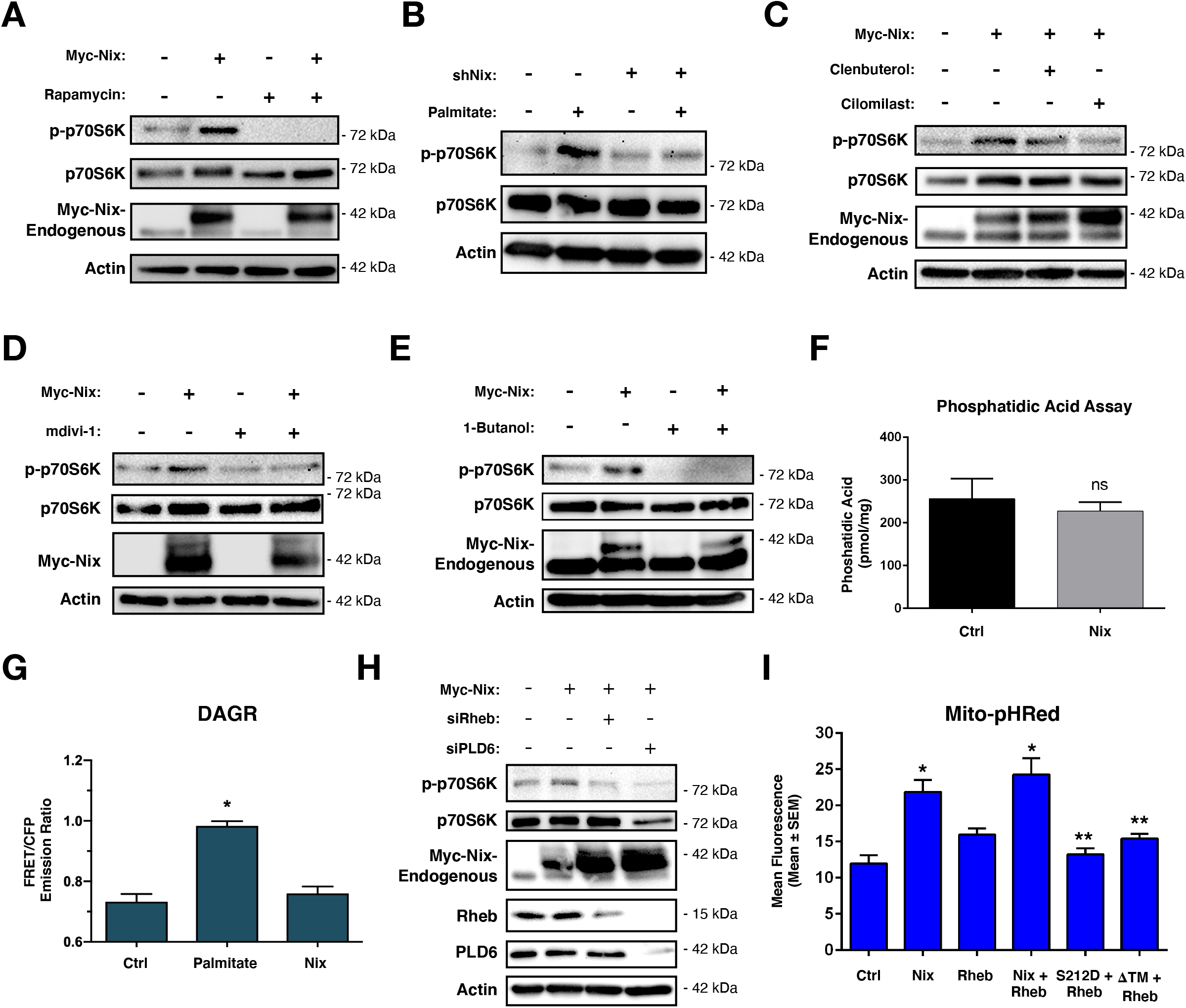
Nix-induced p70S6K activation. A) C2C12 myoblast cells transfected with Myc-Nix or an empty vector control. Cells were treated with Rapamycin (500nM, 1h) or DMSO as vehicle control. Proteins were immunoblotted as indicated. B) C2C12 myoblast cells transfected with shNix, or a scramble control shRNA, and treated overnight with palmitate (200μM) conjugated to 2% albumin in low glucose media. Proteins were immunoblotted as indicated. C) C2C12 myoblast cells were transfected as in (a), and were treated with clenbuterol (500nM), cilomilast (10μM) or vehicle for 2 h. Proteins were immunoblotted as indicated. D) C2C12 myoblast cells were transfected as in (a). Cells were treated with mdivi-1 (20 μM) or vehicle for 1 h. Proteins were immunoblotted as indicated. E) C2C12 myoblast cells were transfected as in (a) and treated with 1-Butanol (1%) for 30 minutes. Proteins were immunoblotted as indicated. F) C2C12 myoblast cells were transfected as in (a), followed by phosphatidic acid assay and quantification. G) C2C12 myoblast cells were transfected as in (a), along with the diacylglycerol biosensor DAGR. Cells were treated overnight with palmitate and analyzed by FRET imaging. H) C2C12 myoblast cells were transfected with Myc-Nix, siRheb, siPLD6. Proteins were immunoblotted as indicated. I) C2C12 myoblast cells were transfected with Myc-Nix, Rheb, Myc-Nix-S212D, Myc-Nix-ΔTM, along with mito-pHRed. Cells were imaged by standard fluorescence and quantified. Data are represented as mean ± S.E.M. *P < 0.05 compared with control, while **P < 0.05 compared with treatment, determined by 1-way or 2-way ANOVA.

As phosphatidic acids have been implicated as important regulators of mTOR signaling in muscle, we used 1-Butanol, a primary alcohol which inhibits phospholipase-D catalyzed phosphatidic acid production, in C2C12 cells expressing Nix. Shown in Figure 6E, Nix-induced p70S6K phosphorylation was completely prevented in the presence of 1-Butanol. To determine if Nix activates mTOR signaling by directly influencing phosphatidic acids production, we expressed Nix in C2C12 cells and evaluated phosphatidic acids content using a commercially available assay. Interestingly, phosphatidic acid content remained unchanged in cells expressing Nix (Fig.6F), as did diacylglycerol signaling (Figure 6G), suggesting that Nix alone does not alter lipid intermediate accumulation in the absence of an ectopic lipid source.

The conversion of phosphatidylcholine into phosphatidic acid involves the enzymatic function of phospholipase D 1 (PLD1), which has been shown to be an important regulator of mTOR activation^32^. However, the mitochondrial-targeted phospholipase D 6 (PLD6) which converts cardiolipin to phosphatidic acid has been shown to modulate mitochondrial dynamics^31^. Moreover, the activation of mTORC1 has been previously shown to be dependent upon lysosomal GTPases, such as Rheb, which can activate mitophagy in a Nix-dependent manner^48^. Thus, we knocked-down both PLD6 and Rheb and determined the effect on Nix-induced p70S6K activation. Shown in Figure 6H, siRNAs targeting Rheb and PLD6 reduced Nix-dependent phosphorylation of p70S6K. In addition, knockdown of PLD6 also reduced the endogenous expression of both total p70S6K and Rheb, suggesting this phospholipase regulates multiple aspects of this pathway. Next, we expressed both Nix and Rheb and observed mito-pHred activation (Figure 6I). The addition of Rheb to Nix enhanced mito-pHred fluorescence; however, when Ser-212 of Nix was mutated to a phospho-mimetic aspartic acid (S212D) or the mitochondrial-targeting transmembrane domain of Nix was deleted (ΔTM), mito-pHred fluorescence was returned to control levels. Collectively, these findings suggest that Nix-induced mitochondrial turn-over modulates Rheb-dependent activation of mTOR-p70S6K, and this pathway has substantial crosstalk with PLD6-dependent production of phosphatidic acid.

## Discussion

Mitochondrial dysfunction has been implicated in insulin resistance and diabetic complications in numerous cell types, although the precise nature of this defect, and consequences resulting from it, are not fully defined in muscle. Utilizing two independent myoblast cell lines, a rodent model, and human iPSC-derived myotubes, we characterized a novel pathway initiated by lipotoxicity resulting in Nix-induced mitochondrial fission, mitophagy, and mTORC1-p70S6K dependent desensitization of IRS1. Interestingly, we also observed crosstalk between phospholipid metabolism and Nix-induced IRS1 inhibition, where phosphatidic acids are critical modulators of this pathway. Furthermore, this work identifies a novel PKA phosphorylation site within the Nix transmembrane domain that serves to translocate Nix from both the mitochondria and ER/SR membranes and inhibit mitochondrial perturbations and p70S6K activation.

Previous work has identified that the calcium-calmodulin dependent phosphatase, calcineurin, activates the mitochondrial fission initiator DRP1^40^. Furthermore, muscle-specific DRP1 deletion in mice results in changes in mitochondrial volume and altered mitochondrial calcium handling^49^. In addition, we have previously demonstrated in cardiomyocytes that Nix modulates ER/SR calcium to control mitochondrial permeability transition^36^. We extend these findings in the present work and demonstrate that Nix-induced ER/SR calcium release activates DRP1 and mitophagy. These observations suggest that in addition to its role as a mitophagy receptor, Nix modulates mitophagy through ER/SR localization and activation of mitochondrial fission. Moreover, in a lipotoxic environment, knockdown of Nix prevents excessive mitophagy, DRP1 activation, and restores insulin-stimulated glucose uptake. Additional evidence also suggests that muscle-specific deletion of the mitophagy receptor Fundc1 protects against HF feeding induced insulin resistance^24^, and that Nix and Fundc1 may cooperative during cardiac progenitor differentiation^50^. Moreover, the Nix homologue Bnip3 is upregulated during myoblast differentiation to protect against oxidative stress, while Bnip3 and p62 are increased in muscle during starvation^51^. Interestingly, we observed that following HF feeding Nix expression was most highly induced in soleus muscle compared to other mitophagy receptors, suggesting it is an important mitophagy regulator during lipotoxic stress. Collectively, these findings suggest that mitophagy receptors may be induced in a stimulus-specific manner to control mitochondrial quality in muscle.

In this report, we also provide detailed mass spectrometry analysis of Nix and identify a novel PKA phosphorylation site within the transmembrane domain. This phospho-acceptor residue serves to inhibit Nix function by promoting the interaction with 14-3-3 chaperones and translocating Nix away from the mitochondria and ER/SR. Using both adrenergic agonists and phosphodiesterase inhibitors, we demonstrate that the function of Nix can be pharmacologically manipulated to modulate myotube mitophagy and insulin sensitivity. These observations are consistent with previous work demonstrating the glucose lowering effects of the phosphodiesterase-4 inhibitor roflumilast^52^ and suggest an additional peripheral mechanism of action of these agents.

One of the most intriguing observations of the present study is the crosstalk between Nix, mitochondrial dynamics, phospholipid metabolism, and mTOR signaling. Phosphatidic acids are a direct activator for mTORC1 both *in vitro* and in cells^53^. Phosphatidic acids bind directly to the FKBP12-rapamycin binding domain of mTOR, and compete with the inhibitor FKBP38 to activate mTORC1^54^. The regulation of mTORC1 also involves recruitment to the lysosomal membranes and activation by small GTPases, such as Rheb, which also compete with FKBP38^55^. Interestingly, stimulation of mitochondrial oxidative phosphorylation results in Nix-dependent recruitment of Rheb to the mitochondria and selective mitochondrial autophagy^(48,56^. Our findings suggest crosstalk between these complex signaling pathways operates during myocyte lipotoxicity, and identifies Nix-induced mitochondrial fission as a regulator of mTOR-p70S6K through Rheb, but contingent on the availability of phosphatidic acids. This notion is consistent with our metabolomics screen, that identified reduced muscle cardiolipin and increased phosphatidic acids content, while our mechanistic cell culture data demonstrate that knockdown of the mitochondrial PLD6 prevents Nix induced p70S6K activation. While the precise relationship between lipotoxicy-induced mitochondrial overload and the regulation of mitochondrial dynamics by phospholipids requires further investigation to more fully define, the activation of mTOR-p70S6K and the phosphorylation of IRS1 at Ser-1101 during lipotoxicity suggests an important mechanism linking excessive muscle mitochondrial turn-over to impaired insulin-stimulated glucose uptake. These observations are consistent with a model whereby Nix responds to lipid-induced mitochondria overload, and protects the myocyte against nutrient storage stress by activating mTOR-p70S6K and IRS1 phosphorylation.

In summary, these studies document a novel signaling cascade triggered by lipotoxicity, and converging on Nix-dependent mitochondrial turn-over and desensitization of insulin receptor signaling. Furthermore, PKA activation downstream of adrenergic signaling inhibits Nix function and restores insulin signaling, suggesting a mechanism by which exercise or pharmacological modulation of PKA may overcome myocyte insulin resistance.

## Materials and Methods

### Plasmids

Myc-tagged Nix, Nix-CytoB5, Nix-MaoB, Nix-ActA, Nix-ΔTM plasmids (Addgene #100795, 100756, 100757, 100758, and 100755) were described previously^36^. The Myc-tagged Nix-S212A and Nix-S212D were generated using a Q5 mutagenesis kit (New England Biolabs). The lentiviral shNix (Addgene #100770) was generated by ligating oligonucleotides containing the targeting sequence 5’-CAGTTCCTGGGTGGAGCTA-3’ into pLKO.1-puro (Addgene # 8453). The mitochondrial (CMV-mitoCAR-GECO1) and endoplasmic reticulum (CMV-ER-LAR-GECO1) targeted calcium biosensors were gifts from Robert Campbell (Addgene #46022 and #61244)^(37,38^. The shNix (#17469)^57^, mEmerald-Mito-7 and mCherry-Mito-7 (#54160, #55102)^58^,^59^, pPHT-PKA (#60936)^60^, DAGR (#14865)^42^, GW1-Mito-pHRed (#31474)^41^ and pcDNA3-FLAG-Rheb (#19996)^61^ plasmids were purchased from Addgene.

### Cell culture and transfections

C2C12 (ATCC CRL-1772) and L6 (ATCC CRL-1458) cell lines were maintained in Dulbecco’s modified Eagle’s medium (DMEM, Hyclone), containing penicillin, streptomycin, and 10% fetal bovine serum (Hyclone) at 37°C and 5% CO_2_. All cells were transfected using JetPrime Polyplus reagent as per the manufacturer’s instructions^35^. The Rheb siRNA (19744) and the Pld6 (194908) were purchased from Dharmacon, and the scrambled siRNA control and control lentivirus were purchased from Santa Cruz (sc-37007, sc-108080). C2C12 and L6 were differentiated by re-feeding confluent cells in 2% fetal bovine serum for to 2–5 days. Human skeletal myoblasts derived from induced pluripotent stem cells were obtained from Cellular Dynamics (iCell SKM-301-020-001-PT). iCell Skeletal Myoblasts were cultured in maintenance medium as per manufacture’s protocol and differentiated for 5-7 days. Palmitate conjugation and treatments were performed as described previously^35^. Clenbuterol, cilomilast, roflumilast, 8-Br-cAMP, forskolin, H89-dihydrochloride hydrate, 2-aminoethoxydipheny borate (2APB), Mdivi-1, etomoxir, rapamycin, and bafilomycin-A1 were purchased from Sigma. Butanol-1 was purchased from Fisher Scientific.

### Immunoblotting and Immunoprecipitation

Protein samples were extracted using a RIPA lysis buffer containing protease and phosphatase inhibitors (Santa Cruz). Subcellular organelle fractionation was performed using a Mitochondrial Isolation Kit (Qiagen Qproteome #37612) and a Nuclear/Cytosolic Isolation Kit (Pierce #78833)^62^. Protein determination was performed using a Bio-Rad protein assay kit and proteins were separated by reducing SDS-PAGE and transferred to a PVDF membrane^35,36,62–64^. Immunoblotting was carried out using the following antibodies for analysis: Bnip3L/Nix (CST #12396), Myc-Tag (CST #2278), HA-Tag (CST #3724), Phospho-p70 S6 Kinase Thr389 (CST #9205), p70 S6 Kinase (CST #9202), Phospho-IRS-1 Ser1101 (CST #2385), IRS1 (ProteinTech #17509-1-AP), Phospho-DRP1 Ser637 (CST #4867), DRP1-D6C7 (CST #8570), Rheb E1G1R (CST #13879), PLD6 (Invitrogen #PA5-71510), BCL2L13 (ProteinTech #16612-1-AP), Parkin (PRK8; CST #4211), BNIP3 (CST #3769), FKBP8 (ThermoFisher #PA5-47513), FUNDC1 (Aviva Systems Biology #ARP53280_P050), AIF (CST #5318), MEK (CST # 8727), SERCA2 (CST #9580), and Actin (Santa Cruz #sc-1616). Antibody dilutions was made as per manufacture’s protocol. To detect the rodent phospho-Bnip3L/Nix, a custom rabbit polyclonal antibody was generated by Abgent using the following peptide sequence IGKRL(pS)TPSAS conjugated to adjuvant. Appropriate horseradish peroxidase-conjugated secondary antibody (Jackson ImmunoResearch Laboratories, USA) was used in a 1:5000 dilution in combination with chemiluminescence to visualize bands using film or a BioRad imager.

### Fluorescent staining

MitoSOX, MitoTracker Red CMXRos, and Hoechst 33342 staining was described previously^35,36,62,65–67^. Glucose uptake assay was measured as previously described using the fluorescent D-glucose analog 2NBDG (200 μM; Molecular Probes)^35^. Microscopy was performed on an Olympus IX70 inverted microscope (Toronto, ON, Canada) with QImaging Retiga SRV Fast 1394 camera (Surrey, BC, Canada) using NIS Elements AR 3.0 software (Nikon Instruments Inc., Melville, NY, USA), or a Zeiss Axiovert 200 inverted microscope fitted with a Calibri 7 LED Light Source and Axiocam 702 mono camera^(36,62^. Quantification, scale bars, background subtract, and processing were done on Fiji (ImageJ) and Zen 2.3 Pro software.

### Transmission electron microscopy (TEM)

TEM imaging was performed according to a protocol described previously^65–67^. Briefly, C2C12 cells myoblasts were seeded in 100 mm plates. Cells were collected using Trypsin. Cells were centrifuged three times (1500×g) and then fixed (3% glutaraldehyde in PBS, pH 7.4) for 3 hours at room temperature. Cells were treated with a post-fixation step using 1% osmium tetroxide in phosphate buffer for 2 hours at room temperature, followed by an alcohol dehydration series before embedding in Epon. TEM was performed with a Philips CM10, at 80 kV, on ultra-thin sections (100 nm on 200 mesh grids). Cells were stained with uranyl acetate and counterstained with lead citrate.

### Phosphatidic acid assay

To quantitatively measure phosphatidic acid (PA) in vitro, a PA assay kit was purchased from PicoProbe™ (BioVision, Milpitas, CA, USA, #K748-100). Protein extractions and fluorometric analysis were performed as per manufacture’s protocol.

### High Fat Diet Animal Model

All procedures in this study were approved by the Animal Care Committee of the University of Manitoba, which adheres to the principles for biomedical research involving animals developed by the Council for International Organizations of Medical Sciences. Male Sprague-Dawley rats were weaned at 3 weeks of age and randomly assigned to a LF diet (10% fat, Research Diets D12450B) or HF diet (45% fat, Research Diets D12451), both containing sucrose, for 12 weeks, as previously described^(35,68^. Metabolomics analysis was performed on a 1290 Infinity Agilent high-performance liquid chromatography (HPLC) system coupled to a 6538 UHD Agilent Accurate Q-TOF LC/MS equipped with a dual electrospray ionization source, as previously described^(35,68^. PCR-based gene expression arrays were purchased from SA Biosciences (Qiagen).

### In vitro kinase assay and Phospho-peptide mapping

Synthetic peptides (GeneScript) were resuspended at a concentration of 1 mg/ml. These peptides were used as the substrate in a PKA kinase assay kit (New England Biolabs, #P6000S) according to the manufacturer’s instructions, with the exception that [32P]-ATP was replaced with fresh molecular biology grade ATP. The Kemptide substrate (Enzo Life Sciences; #P-107; LRRASLG) was used as a positive control in each assay. Before mass spectrometry analysis, kinase assays were prepared using C18 ZipTips (EMD Millipore, Etobicoke, ON, Canada). Samples in 50% acetonitrile and 0.1% formic acid were introduced into a linear ion-trap mass spectrometer (LTQ XL: ThermoFisher, San Jose, CA, USA) via static nanoflow, using a glass capillary emitter (PicoTip: New Objective, Woburn, MA, USA), as described previously^35^.

### Statistics

Data are presented as mean ± standard error of the mean (S.E.M.). Differences between groups in imaging experiments with only 2 conditions were analyzed using an unpaired t-tests, where * indicates P < 0.05 compared with control. Experiments with 4 or more conditions were analyzed using a 1-way ANOVA or 2-way ANOVA, with Tukey’s test for multiple comparison, where * indicates P < 0.05 compared with control, and ** indicates P < 0.05 compared with treatment. All statistical analysis were performed using GraphPad Prism 6 software.

## Acknowledgments

This work was support by the Natural Science and Engineering Research Council (NSERC) Canada, through Discovery Grants to JWG and ARW. Seed funding was provided by the Children’s Hospital Research Institute of Manitoba, the DREAM research theme, and the Manitoba Centre for Nursing and Health Research. V.W.D is supported by CIHR and is the Allen Rouse Basic Scientist of the Manitoba Medical Services Foundation. J.W.G., A.R.W., and V.W.D are members of the DEVOTION Research Cluster. S.C.d.S.R. is supported by a University of Manitoba Graduate Studentship, J.T.F. is supported by an Alexander Graham Bell studentship from NSERC Canada, and M.D.M. and S.M.K. are supported by a studentship from the Children’s Hospital Foundation of Manitoba and Research Manitoba.

