## Supplemental Data for "Nix induced mitochondrial fission, mitophagy, and myocyte insulin resistance are abrogated by PKA phosphorylation"

Supplemental Figure 1

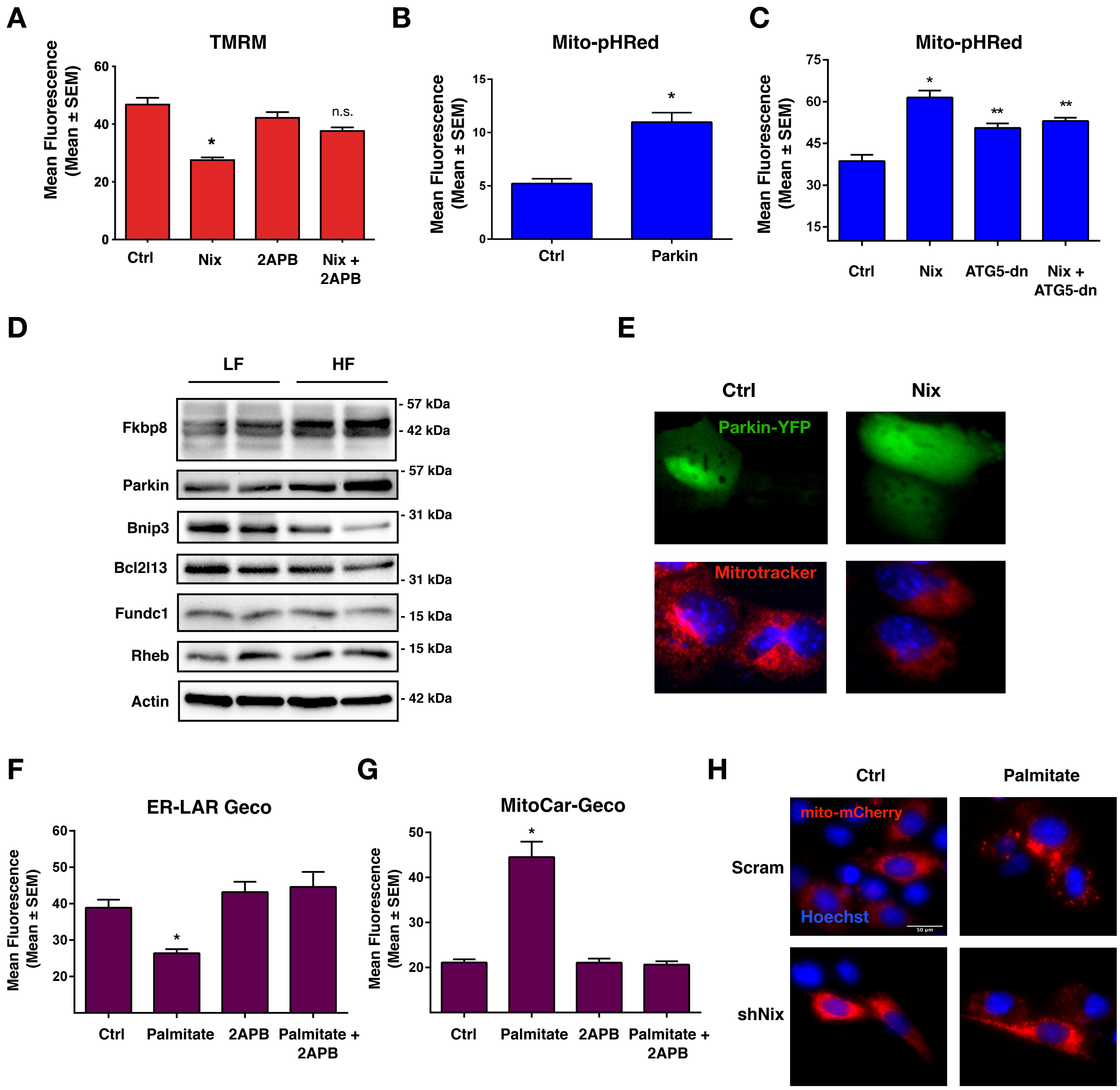

Supplemental Figure 2

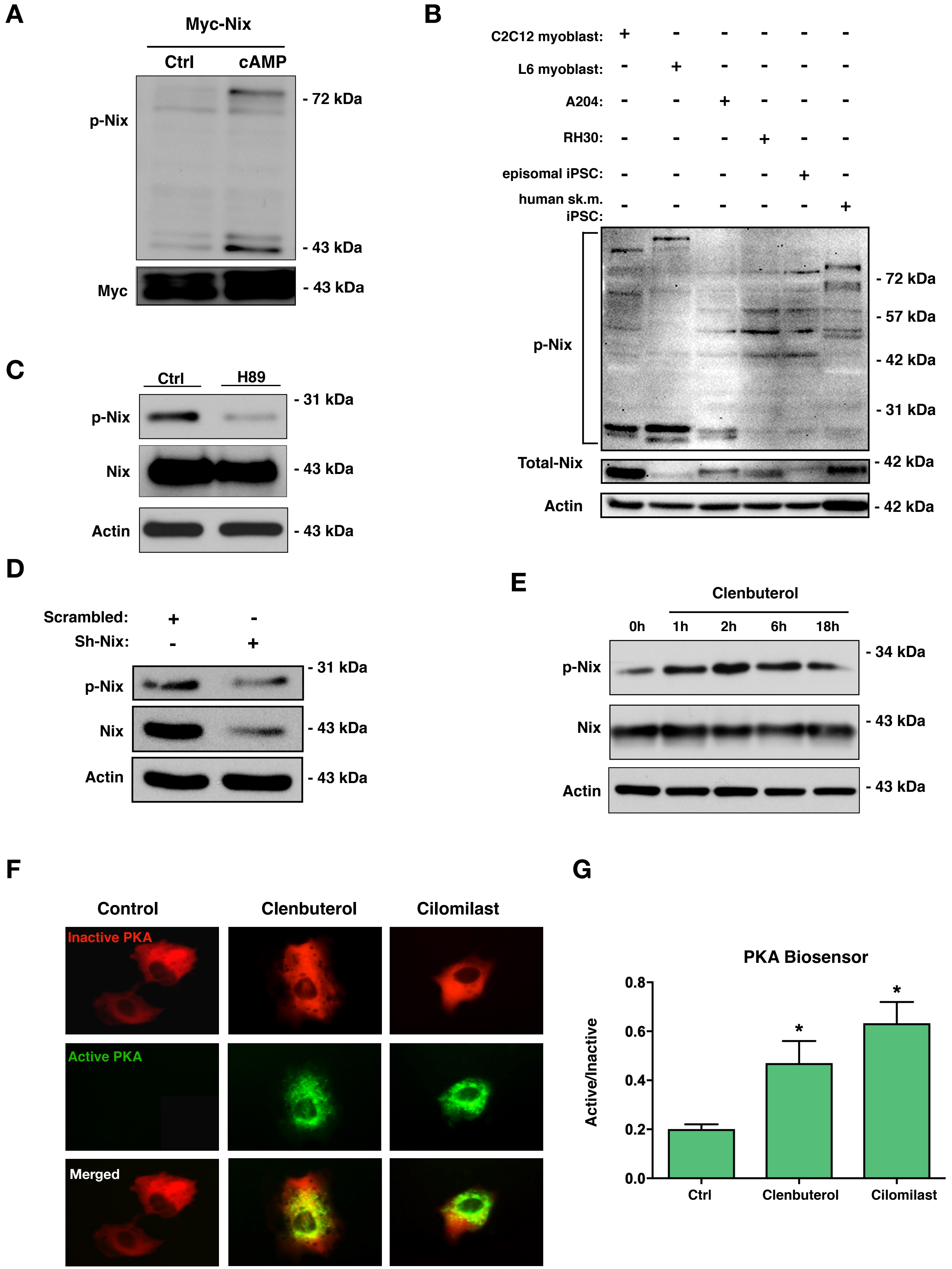

Supplemental Figure 3.

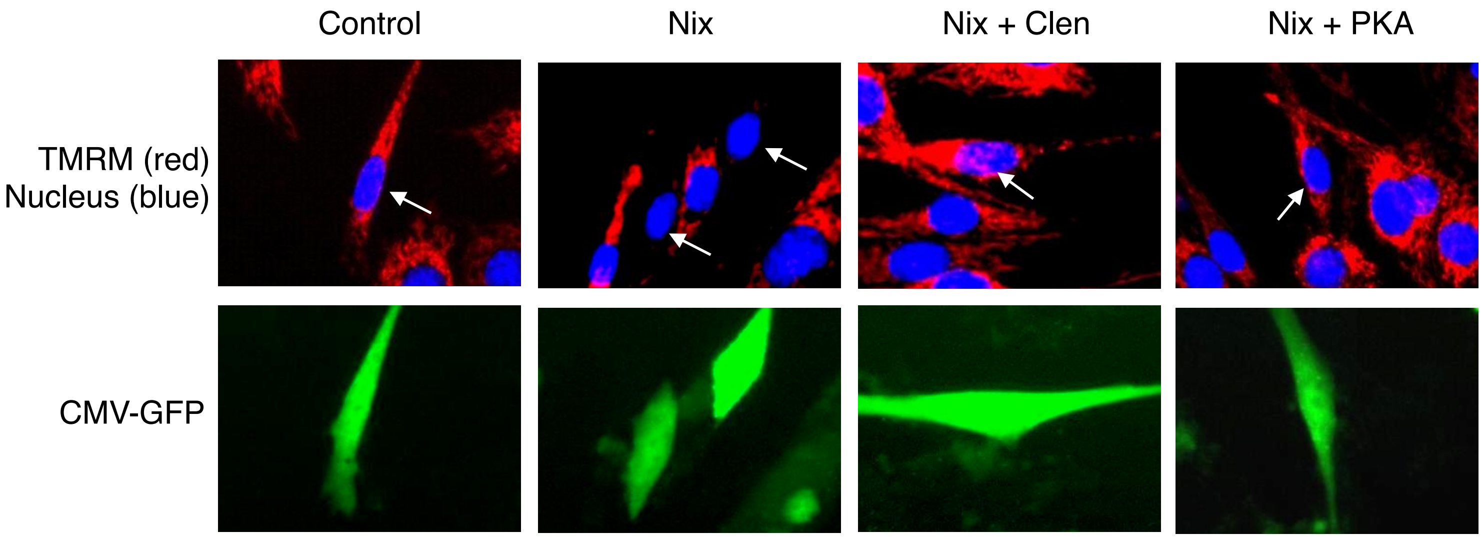

**Supplemental Figure 1.** ***Palmitate-induced mitochondrial dysfunction.***

A) C2C12 myoblast cells were transfected with Myc-Nix or an empty vector. Cells were treated with 2-aminoethoxydiphenyl borate (2APB 10µM, 1h), or DMSO as a vehicle control. Cells were stained for TMRM and quantified. B) C2C12 myoblasts cells were transfected with HA-Parkin or an empty vector, along with the mitophagy biosensor mito-pHRed. C) C2C12 myoblast cells were transfected Myc-Nix, a dominant-negative ATG5, or an empty vector, along with mito-pHRed. D) Western blot analysis of rat soleus muscle (n=2) exposed to high fat (HF) or low fat (LF) diet for 12-weeks. Protein extracts were analyzed as indicated. E) C2C12 myoblasts cells were transfected with Myc-Nix or an empty vector, along with fluorescent protein YFP-Parkin. Cells were stained with Mitotracker and Hoechst and imaged by standard fluorescence microscopy. F-G) C2C12 myoblasts cells were transfected with ER-Lar-Geco (f) or MitoCar-Geco (g) and an empty vector, followed by overnight treatment with palmitate conjugated to 2% albumin in low glucose media. Control cells were treated with 2% albumin alone. Following palmitate treatment, cells were treated with 2APB, as described in (a). H) C2C12 myoblasts were transfected with shNix, or a scrambled control shRNA, along with mito-mCherry. Cells were treated overnight with palmitate conjugated to 2% albumin in low glucose media. Control cells were treated with 2% albumin alone. Cells were stained with Hoechst and imaged by standard fluorescence microscopy. Data are represented as mean ± S.E.M. *P < 0.05 compared with control, while **P < 0.05 compared with treatment, determined by 1-way ANOVA.

**Supplemental Figure 2.** ***PKA Phosphorylation of Nix.***

A) C2C12 myoblast cells were transfected with Myc-Nix or an empty vector and treated with c-AMP analogue (1mM, 1h) or vehicle control. Protein extracts were immunoblotted, as indicated. B) Protein extract of multiple cell lines (as indicated) were immunoblotted for endogenous phospho-Nix. C) C2C12 myoblast cells were treated with PKA inhibitor, H89 (10 µM, 1h) or vehicle control. Protein extracts were immunoblotted, as indicated. D) C2C12 myoblasts were transfected with shNix, or a scrambled control shRNA. Protein extracts were immunoblotted, as indicated. E) C2C12 myoblast cells were treated with 500 nM clenbuterol or vehicle for multiple time points. Protein extracts were immunoblotted, as indicated. F) C2C12 myoblast cells were transfected with PKA biosensor (pPHT-PKA), and treated with clenbuterol (500nM), cilomilast (10µM) or vehicle for 2 h. Cells were imaged by standard fluorescence microscopy. G) Quantification of fluorescent images in (f) by measuring the ratio of green (active) to red (inactive) fluorescent signal. Data are represented as mean ± S.E.M. *P < 0.05 compared with control, determined by 1-way ANOVA.

**Supplemental Figure 3.** ***Nix-induced mitochondrial depolarization.***

A) C2C12 myoblast cells were transfected with Nix, PKA or CMV-GFP as a control. Cells were treated with clenbuterol (500nM, 2h) or a vehicle control. Cells were stained for TMRM and Hoechst and imaged by standard fluorescence. Red active or inactive mitochondria are indicated by arrows.

### Table 1: Metabolomics Table for Selected Muscle Triglycerides

| TG | HF diet (Fold of Control) |
|---|---|
| TG(22:3/22:6/22:6) | 474.16094524 |
| TG(21:0/22:0/22:3) | 225.30109372 |
| TG(20:0/22:0/22:6) | 252.24362556 |
| TG(13:0/18:3/22:2) | 0.73820095 |
| TG(13:0/16:0/18:3) | 0.00002156 |
| TG(13:0/15:1/17:2) | 12.42594972 |
| TG(13:0/14:0/18:2) | 15.80467647 |
| TG(13:0/14:0/18:2) | 0.03724308 |
| TG(12:0/19:1/19:1) | 0.00010212 |
| TG(12:0/17:0/17:0) | 0.00000017 |
| TG(12:0/16:0/20:1) | 0.00454675 |
| TG(12:0/15:1/16:0) | 0.09477939 |
| TG(12:0/15:1/16:0) | 226.25782085 |
| TG(12:0/14:1/18:4) | 451.47187440 |
| TG(12:0/14:1/18:0) | 0.03885087 |
| TG(12:0/14:1/15:1) | 303.10081212 |
| TG(12:0/14:1/15:1) | 5701844.90556225 |
| TG(12:0/13:0/22:5) | 27.28222775 |
| TG(12:0/12:0/20:0) | 215.98445754 |
| TG(12:0/12:0/19:1) | 14.43848324 |
| TG(12:0/12:0/18:0) | 16.99116745 |
| TG(12:0/12:0/14:1) | 378.78275633 |

### Table 2: Metabolomics Table for Selected Muscle Diacylglycerides

| DG | HF diet (Fold of Control) |
|---|---|
| DG(O-16:0/18:1) | 302.8633348 |
| DG(22:3/22:6) | 315.3130896 |
| DG(22:3/22:2) | 0.765006279 |
| DG(22:2/24:1) | 685.4931195 |
| DG(22:2/24:0) | 393.766622 |
| DG(21:0/22:6) | 271.0025641 |
| DG(20:5/22:4) | 380753.4087 |
| DG(20:4/22:6) | 6176.598903 |
| DG(20:4/20:4) | 16.2638058 |
| DG(20:2/24:0) | 0.021997727 |
| DG(20:2/22:0) | 0.643402435 |
| DG(20:2/14:1) | 244.0439805 |
| DG(20:2/20:2) | 16.8409959 |
| DG(20:1/18:0) | 388.6799158 |
| DG(20:0/22:1) | 208.6105653 |
| DG(20:0/20:0) | 0.3638602 |
| DG(19:0/19:0) | 12.66929387 |
| DG(18:4/24:1) | 255.600736 |
| DG(18:4/22:6) | 499.6432217 |
| DG(18:2/18:1) | 19.57482869 |
| DG(18:1/24:1) | 0.040373546 |
| DG(17:0/17:0) | 0.00292789 |
| DG(16:1/16:1) | 347.202625 |
| DG(16:0/20:0) | 306.2108676 |
| DG(16:0/18:0) | 8408.447173 |
| DG(16:0/16:0) | 346.1952462 |
| DG(15:0/16:0) | 0.046122772 |
| DG(12:0/20:0) | 9857.486705 |

### Table 3: Metabolomics Table for Selected Muscle Cerimides

| Cer | HF diet (Fold of Control) |
|---|---|
| α-Galactosyl Ceramide | 0.002657066 |
| Lactosylceramide (d18:1/22:0) | 16.50345981 |
| GlcCer(d18:2/23:0) | 7.268878339 |
| GlcCer(d18:2/23:0) | 3059839.875 |
| GlcCer(d18:2/23:0) | 2728569.373 |
| GlcCer(d18:2/21:0) | 557.2568299 |
| GlcCer(d18:2/21:0) | 12.48381689 |
| GlcCer(d18:1/26:0) | 4.72854E-07 |
| GlcCer(d18:0/24:0) | 3.28482E-06 |
| GlcCer(d16:2/20:0) | 19.35893404 |
| GlcCer(d15:2/22:0) | 0.409136853 |
| GlcCer(d15:2/22:0) | 328.6126366 |
| GlcCer(d15:2/22:0) | 34.65256394 |
| GlcCer(d15:2/20:0) | 8118.391671 |
| GlcCer(d15:2/18:0) | 293.9558449 |
| GlcCer(d15:1/22:0) | 15.98534449 |
| GlcCer(d15:1/20:0) | 371.4533676 |
| GlcCer(d15:1/20:0) | 324.6849811 |
| GlcCer(d14:1/18:1) | 348.5725632 |
| Dihydroceramide C2 | 556.0446801 |
| Ceramide (d18:1/26:0) | 205.005966 |
| Cer(t18:0/16:0) | 5829.65025 |
| Cer(t18:0/16:0) | 14.57712641 |
| Cer(t18:0/16:0) | 216.174369 |
| Cer(d18:2/23:0) | 0.030951015 |
| Cer(d18:2/20:1) | 270.0500114 |
| Cer(d18:2/20:1) | 297.7411079 |
| Cer(d18:2/20:1) | 11.30894339 |
| Cer(d18:2/18:1) | 0.944696921 |
| Cer(d18:2/14:0) | 16.71667118 |
| Cer(d18:2/14:0) | 282.8826293 |
| Cer(d18:1/22:1) | 0.020334332 |
| Cer(d18:1/20:0) | 2.15693E-06 |
| Cer(d18:1/16:0) | 367.3849827 |
| Cer(d18:1/16:0) | 190.163429 |
| Cer(d18:0/17:0) | 225.1782813 |
| Cer(d18:0/15:0) | 828.6573179 |
| Cer(d18:0/13:0) | 6573251.966 |
| Cer(d18:0/12:0) | 117007904.1 |
| Cer(d16:2/24:0) | 213.6848341 |
| Cer(d16:2/20:1) | 122962.2496 |
| Cer(d16:1/23:0) | 271.2296103 |

| | |
|---|---|
| Cer(d15:2/22:0) | 0.803124732 |
| Cer(d14:2/20:0) | 423.0099666 |
| Cer(d14:2/18:1) | 0.028376394 |
| Cer(d14:2/18:0) | 10.79612912 |
| Cer(d14:2/18:0) | 1.51852E+11 |
| Cer(d14:1/26:0) | 450314.2601 |
| Cer(d14:1/24:0) | 15.3688333 |
| Cer(d14:1/22:1) | 298.2445388 |
| Cer(d14:1/22:0) | 509.2305146 |
| C-6 NBD Ceramide | 450.8602007 |
| AV-Ceramide | 283.9401451 |
| PE-Cer(d16:1/23:0) | 13621.34993 |
| PE-Cer(d16:1/18:0) | 14.52539596 |
| PE-Cer(d14:1/25:0) | 344.1520018 |
| PE-Cer(d14:1/23:0) | 10237.21749 |
| CerP(d18:1/24:1) | 140907.4644 |
| CerP(d18:1/18:0) | 273.3677637 |

### Table 4: Gene expression Array Table for Selected Muscle Cardiolipins

| CL | HF diet (Fold of Control) |
|---|---|
| CL(18:2/18:2/18:2/18:2) | 0.002684369 |
| CL(22:6/20:3/18:2/18:2) | ND in HF condition |
| CL(18:2/18:2/18:2/18:3) | ND in HF condition |
| CL(22:1/22:1/22:1/14:1) | ND in HF condition |
| CL(14:1/14:1/14:1/15:1) | ND in HF condition |

**Table 5: Gene expression Array Table for Selected Muscle Phosphatidic Acids**

| PA | HF diet (Fold of Control) |
|---|---|
| PA(20:0/22:6) | 19842.77007 |
| PA(20:0/18:2) | 509.6983175 |
| PA(22:6/14:1) | 0.05364954 |
| PA(20:5/22:6) | 0.000120273 |
| PA(20:5/18:3) | 245.1649605 |
| PA(20:5/18:3) | 13.27455068 |
| PA(20:5/18:3) | 0.526316154 |
| PA(20:5/18:3) | 300.8537298 |
| PA(20:3/21:0) | 518.1851107 |
| PA(20:2/21:0) | 296.0700657 |
| PA(20:1/22:0) | 243.7446545 |
| PA(19:0/20:0) | 12388.91034 |
| PA(18:4/20:5) | 12.8543701 |
| PA(18:4/18:4) | 314.6167207 |
| PA(18:4/18:3) | 0.072812107 |
| PA(16:0/21:0) | 366.4071164 |
| PA(15:1/22:4) | 335.3509184 |
| PA(15:0/20:3) | 206.8535602 |
| PA(14:1/17:2) | 0.708617217 |
| PA(14:0/15:0) | 4534.707717 |
| PA(14:0/12:0) | 257.1628321 |

### Table 6: Gene expression Array Table for Selected Muscle Genes

| Gene | HF diet (Fold of Control) |
|---|---|
| Atg12 | 0.82 |
| Atg16l1 | 0.97 |
| Atg3 | 0.86 |
| Atg5 | 0.73 |
| Atg7 | 1.13 |
| Atp6v1g2 | 0.39 |
| Bax | 1.05 |
| Bcl2 | 1.08 |
| Bcl2a1 | 0.7 |
| Bcl2l1 | 0.7 |
| Bcl2l11 | 0.91 |
| Becn1 | 0.84 |
| Igf1 | 1.13 |
| Igf1r | 1.25 |
| Tnf | 0.69 |
| Tnfrsf11b | 0.69 |
| Tnfrsf1a | 0.94 |
| Tp53 | 1.32 |
| Ulk1 | 1.03 |
| Actb | 1.08 |
| Ldha | 1.03 |
| Cs | 0.8 |
| Foxo3 | 0.87 |
| Hdac5 | 0.91 |
| Mb | 0.48 |
| Mef2c | 0.91 |
| Mstn | 1.51 |
| Myf5 | 1.37 |
| Myod1 | 1.25 |
| Myog | 0.89 |
| Myh1 | 0.38 |
| Myh2 | 0.04 |
| Nfkb1 | 1.04 |
| Pdk4 | 0.51 |
| Pparg | 0.85 |
| Ppargc1a | 0.65 |
| Ppargc1b | 0.98 |
| Rhoa | 0.97 |
| Slc2a4 | 0.95 |
| Tgfb1 | 1.19 |
| Tnnc1 | 0.02 |
| Tnni2 | 1.02 |

| | |
|---|---|
| Tnnt1 | 0.02 |
| Tnnt3 | 1.06 |
| Fasn | 0.18 |
| Gys1 | 0.95 |
| Hk2 | 0.8 |
| Il6 | 0.58 |
| Insr | 0.89 |
| Irs1 | 1.2 |
| Irs2 | 0.96 |
| Mtor | 0.97 |
| Slc27a1 | 0.66 |
| Srebf1 | 1.14 |
| Srebf2 | 1.17 |
| Bnip3 | 0.84 |
| Cox18 | 0.91 |
| Cpt1b | 0.82 |
| Cpt2 | 0.78 |
| Dnm1l | 0.73 |
| Fis1 | 0.82 |
| Gpx1 | 0.51 |
| Mfn1 | 0.74 |
| Mfn2 | 0.86 |
| Opa1 | 0.92 |
| Taz | 0.91 |
| Timm44 | 0.96 |
| Tomm40 | 1.02 |
| Tomm22 | 0.94 |
| Ucp2 | 1.11 |
| Ucp3 | 0.8 |
| Acly | 0.48 |
| Aldoa | 1.01 |
| Idh2 | 0.74 |
| Mdh1 | 0.63 |
| Mdh2 | 0.91 |
